# The effect of strong purifying selection on genetic diversity

**DOI:** 10.1101/211557

**Authors:** Ivana Cvijović, Benjamin H. Good, Michael M. Desai

## Abstract

Purifying selection reduces genetic diversity, both at sites under direct selection and at linked neutral sites. This process, known as background selection, is thought to play an important role in shaping genomic diversity in natural populations. Yet despite its importance, the effects of background selection are not fully understood. Previous theoretical analyses of this process have taken a backwards-time approach based on the structured coalescent. While they provide some insight, these methods are either limited to very small samples or are computationally prohibitive. Here, we present a new forward-time analysis of the trajectories of both neutral and deleterious mutations at a nonrecombining locus. We find that strong purifying selection leads to remarkably rich dynamics: neutral mutations can exhibit sweep-like behavior, and deleterious mutations can reach substantial frequencies even when they are guaranteed to eventually go extinct. Our analysis of these dynamics allows us to calculate analytical expressions for the full site frequency spectrum. We find that whenever background selection is strong enough to lead to a reduction in genetic diversity, it also results in substantial distortions to the site frequency spectrum, which can mimic the effects of population expansions or positive selection. Because these distortions are most pronounced in the low and high frequency ends of the spectrum, they become particularly important in larger samples, but may have small effects in smaller samples. We also apply our forward-time framework to calculate other quantities, such as the ultimate fates of polymorphisms or the fitnesses of their ancestral backgrounds.

## INTRODUCTION

Purifying selection against newly arising deleterious mutations is essential to preserving biological function. This process is ubiquitous across all natural populations, and is responsible for genomic sequence conservation across long evolutionary timescales. In addition to preserving function at directly selected sites, negative selection also leaves signatures in patterns of diversity at linked neutral sites that have been observed in a wide range of organisms (Begun and Aquadro, 1992; Charlesworth, 1996; Comeron, 2014; Cutter and Payseur, 2003; Elyashiv et al., 2016; Flowers et al., 2012; McVicker et al., 2009). This process is known as background selection, and understanding its effects is essential to characterizing the evolutionary pressures that have shaped a population, as well as to distinguishing its effects from less ubiquitous events such as population expansions or the positive selection of new adaptive traits.

At a qualitative level, the effects of background selection are well-known: it reduces linked neutral diversity by reducing the number of individuals that are able to contribute descendants in the long run. Since individuals that carry strongly deleterious mutations cannot leave descendants on long timescales, all diversity that persists in the population must have arisen in individuals that were free of deleterious mutations. Since all of these individuals are equivalent in fitness, this suggests that diversity should resemble that expected in a neutral population of smaller size — specifically, with size equal to the number of mutation-free individuals (Charlesworth et al., 1993).

However, an extensive body of work has shown that this intuition is not correct, and that background selection against strongly deleterious mutations can lead to non-neutral distortions in diversity statistics (Charlesworth et al., 1993, 1995; Good et al., 2014; Gordo et al., 2002; Hudson and Kaplan, 1994; Nicolaisen and Desai, 2012; O’Fallon et al., 2010; Tachida, 2000; Walczak et al., 2011; Williamson and Orive, 2002). The reason for this is simple: even strong selection cannot purge deleterious alleles instantly. Instead, deleterious haplotypes persist in the population on short timescales, allowing neutral variants that arise on their backgrounds to reach modest frequencies. This is most readily apparent in statistics based on the site frequency spectrum (the number, *p*(*f*), of polymorphisms which are at frequency *f* in the population), such as the number of singletons or Tajima’s D (Tajima, 1989). As we show below, even when deleterious mutations have a strong effect on fitness, the site frequency spectrum shows an enormous excess of rare variants compared to the expectation for a neutral population of reduced effective size.

These signatures in genetic diversity are qualitatively similar to those we expect from population expansions and positive selection (Keinan and Clark, 2012; Rannala, 1997; Sawyer and Hartl, 1992; Slatkin and Hudson, 1991). A detailed quantitative understanding of background selection is therefore essential if we are to disentangle its signatures from those of other evolutionary processes.

The traditional approach to analyzing the effects of purifying selection has been to use backwards-time approaches based on the structured coalescent (Hudson and Kaplan, 1994, 1988). This offers a framework to model how background selection affects the statistics of genealogical histories of a sample, and hence the expected patterns of genetic diversity. However, while these backwards-time structured coalescent methods make it possible to rapidly simulate genealogies, they are essentially numerical methods and do not lead to analytical predictions. A more technical but crucial limitation is that they rapidly become intractable in larger samples. This is becoming an increasingly important problem as advances in sequencing technology now make it possible to study sample sizes of thousands (or even hundreds of thousands) of individuals. The poor scaling of coalescent methods with sample size is of particular importance in studying background selection: since purifying selection is expected to result in an excess of rare variants, its effects increase in magnitude as sample size increases. This can reveal deviations from neutrality in large samples that are not seen in smaller samples.

Here, we use an alternative, forward-time approach to analyze how purifying selection affects patterns of genetic variation. Our method is based on the observation that to predict single-locus statistics, such as the site frequency-spectrum, it is not necessary to model the entire genealogy. Instead, we model the frequency of the lineage descended from a single mutation as it changes over time due to the combined forces of selection and genetic drift, and as it accumulates additional deleterious mutations. We then use these allele frequency trajectories to predict the site frequency spectrum, from which any other single-site statistic of interest can then be calculated.

We show that background selection creates large distortions in the frequency spectrum at linked neutral sites whenever there is significant fitness variation in the population. These distortions are concentrated in the high and low frequency ends of the frequency spectrum, and hence are particularly important in large samples. We provide analytical expression for the frequencies at which these distortions occur, and can therefore predict at what sample sizes they can be seen in data.

Aside from single-timepoint statistics such as the site frequency spectrum, we also obtain analytical forms for the statistics of allele frequency trajectories. These trajectories have a very non-neutral character, which reflects the underlying linked selection. Our approach offers an intuitive explanation for how these non-neutral behaviors arise in the presence of substantial linked fitness variation, which explains the origins of the distortions in the site frequency spectrum.

The statistics of allele frequency trajectories can also be used to calculate any time-dependent single-site statistic. For example, we analyze how the future trajectory of a mutation can be predicted from the frequency at which we initially observe it, and we discuss the extent to which the the observed frequency of a polymorphism can inform us about the fitness of the background on which it arose.

We begin in the next section by providing an intuitive explanation for the origins of the distortions in the site frequency spectrum in the presence of strong background selection, and explain why these distortions always accompany a reduction in diversity. This section summarizes the importance of correctly accounting for background selection, particularly when analyzing large samples, and should be accessible to all readers. We next explain how dynamical aspects of allele frequency trajectories can be related to site frequency spectra, using the trajectories of unlinked loci as an example. Readers already familiar with this intuition may choose to skip ahead to the background selection model, but those less interested in the technical details may find that this section provides useful intuition for the calculations in a simpler context.

We next define a specific model of background selection and present an intuitive description of the key features of allele frequency trajectories. These sections are also largely non-technical, and may be of interest to readers who wish to understand the origins of non-neutral behaviors of alleles in the presence of strong background selection. Finally, in the Analysis, we turn to a formal stochastic treatment of the trajectories of neutral and deleterious mutations. In the last section, we use these trajectories to calculate the site frequency spectrum and other statistics describing genetic diversity within the population.

## STRONG BACKGROUND SELECTION DISTORTS THE SITE FREQUENCY SPECTRUM

We begin by presenting a more detailed intuitive description of the effects of background selection on linked neutral alleles. We focus on analyzing the allele frequency spectrum, defined as the expected number, *p*(*f*), of mutations that are present at frequency *f* within the population in steady state. This allele frequency spectrum contains all relevant information about single-site statistics: any such statistic of interest can be calculated by subsampling appropriately from *p*(*f*).

In Figure 1A, we show an example of the site frequency spectrum of neutral mutations at a locus experiencing strong background selection, generated by Wright-Fisher forward time simulations. This example shows several key generic features of background selection. First, at intermediate frequencies the site frequency spectrum has a neutral shape, *p*(*f*) ∝ *f*^−1^, with the total number of such intermediate-frequency polymorphisms consistent with the simple reduced effective population size prediction (Charlesworth et al., 1993). However, at both low and high frequencies *p*(*f*) is significantly distorted. At low frequencies, we see an enormous excess of rare alleles, qualitatively similar to what we expect in expanding populations (Ran-nala, 1997; Slatkin and Hudson, 1991). We also see a large excess of very high frequency variants, leading to a non-monotonic site frequency spectrum. This is reminiscent of the non-monotonicity seen in the presence of positive selection (Sawyer and Hartl, 1992). Notably, these distortions at both high and low frequencies arise in populations of constant size in which all variation is either neutral or deleterious.

**FIG. 1.**
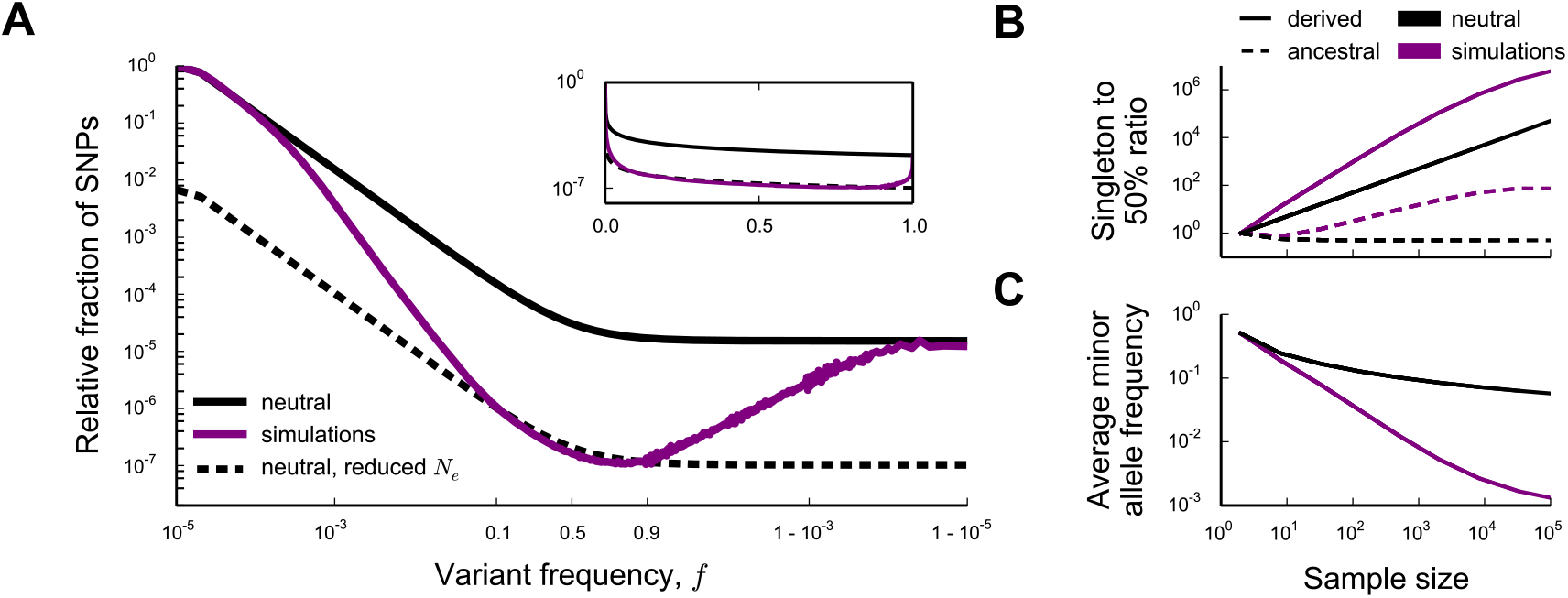
(A) The (unfolded) average site frequency spectrum of neutral alleles along a nonrecombining genomic segment in an asexual population of N = 10^5^ individuals deviates strongly from the prediction of neutral theory. Deleterious mutations occur at rate *NU*_*d*_ = 5000 and all have the same effect on fitness *N*_*s*_ = 1000 (purple). The black and grey lines show the neutral expectation for the site frequency spectrum of a population of 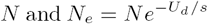 individuals, respectively. The inset shows the same data, but with the x-axis linearly scaled to emphasize intermediate frequencies. Averages were generated by bootstrap resampling with replacement of mutations that arose in 10^5^ Wright-Fisher simulations in which neutral mutations were set to occur at rate *NU*_*n*_ = 10^3^, and by smoothing the obtained curve using a box kernel smoother of width much smaller than the scale on which the site frequency spectrum varies (the kernel width was set to < 10% of the minor allele frequency). (B, C) Statistics of the site frequency spectrum in samples of the population shown on the left (purple) can deviate from predictions of neutral theory (black) by many orders of magnitude in large samples, even though the effect of background selection will be small in small samples. Here shown: (B) The average ratio of the number polymorphisms present as derived singletons (full lines) or ancestral singletons (dotted lines) in the sample to the number present at 50% frequency, and (C) the average minor allele frequency of the sampled alleles.

The excess of rare derived alleles arises because selection takes a finite amount of time to purge deleterious genotypes. Thus we expect that there can be substantial neutral variation linked to deleterious alleles that, though doomed to be eventually purged from the population, can still reach modest frequencies. At the very lowest frequencies, we expect that neutral mutations arising in all individuals in the population (independent of the number of deleterious mutations they carry) can contribute. Thus, at the lowest frequencies, the site frequency spectrum should be unaffected by selection, and should agree with the neutral site frequency spectrum of a population of size *N*. On the other hand, as argued above, the total number of common alleles must reflect the (much smaller) number of deleterious mutation free individuals, because only neutral mutations arising in such individuals can reach such high frequencies. Since the overall number of very rare alleles is proportional to the census population size *N*, and the number of common alleles reflects a much smaller deleterious mutation-free subpopulation, there must be a transition between these two: between these extremes the site frequency spectrum must fall off more rapidly than the neutral prediction *p*(*f*) ~ *f*^−1^. This transition reflects the fact that as frequency increases, the effect of selection will be more strongly felt, and neutral mutations arising in genotypes of increasingly lower fitnesses will become increasingly unlikely.

As the frequency increases even further, we see from our simulations that the total number of polymporphisms increases again, until at very high frequencies it matches the prediction for a neutral population of size equal to the census size *N.* Note that at these frequencies, the total number of backgrounds contributing to the diversity is constant (i.e. all mutations reaching these frequencies must arise in the small subpopulation of mutation-free individuals). This suggests that fundamentally non-neutral behaviors must be dominating the dynamics of these high frequency neutral polymorphisms. To understand this, as well as the details of the rapid fall-off at very low frequencies, we will need to develop a more detailed description of the trajectories of neutral alleles in the population; we analyze this in quantitative detail in a later section.

However, a simple argument can explain the agreement with the neutral prediction at the highest frequencies. Polymorphisms observed at these very high frequencies correspond to neutral variants that have almost reached fixation. The ancestral allele is still present in the population, but at a very low frequency. In principle, the dynamics of the derived and ancestral alleles should depend on the fitnesses of their backgrounds. However, once the frequency of the ancestral allele is sufficiently low, selection is no longer effective and its dynamics must become neutral, independent of what it is linked to. At steady state, the total rate at which neutral mutations fix is equal to the product of the rate at which they enter the population at any point in time (*NU*_*n*_) and their fixation probability, 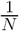, (Birky and Walsh, 1988). Thus, the overall rate at which neutral mutations enter this high frequency regime agrees with the rate in a neutral population at the census population size. Since the total rate at which alleles enter this high frequency regime is unaffected by selection, and since their dynamics within this regime are neutral, we expect that the site-frequency spectrum should also agree with the neutral prediction for a population of size *N.*

Although these simple arguments do not provide a full quantitative explanation of the site frequency spectrum, they already offer some intuition about the presence and magnitude of the distortions due to background selection. First, these distortions arise in part as a result of the difference in the number of backgrounds on which mutations that remain at the lowest frequencies and mutations that reach substantial frequencies can arise. Thus, they will always occur when background selection is strong enough to cause a substantial reduction in the “effective population size”: if the pairwise diversity π is at all reduced compared to the neutral expectation 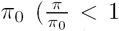, or, in terms of McVicker’s B-statistic, *B* < 1, (McVicker et al., 2009)), these distortions exist (see Figure 1A). Second, because the distortions from the neutral shape are limited to high and low ends of the frequency spectrum, they will have limited effect on site frequency spectra of small samples, but will have dramatic consequences as the sample size increases (see Figure 1B,C). On a practical level, this means that extrapolating conclusions from small samples about the effects of background selection can be grossly misleading.

In the next few sections, we will analyze the dynamics of neutral mutations under background selection in detail. However, before analyzing a model of background selection, we begin in the next section by reviewing a heuristic argument explaining the intuition for the allele frequency trajectories and the shape of the site-frequency spectrum of single, unlinked loci.

## UNLINKED LOCI

To gain insight into the more complicated case of linked selection, we first begin by reviewing the simplest case of a single locus unlinked to any other selected loci. The probability that an allele at that locus is present at frequency *f* at time *t*, *p*(*f*, *t*), is described by the diffusion equation

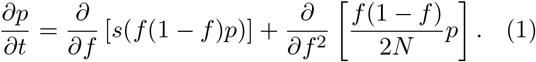

Ewens (1963) showed that the expected site frequency spectrum (SFS) can be obtained from this forward-time description of the allele frequency trajectory: because mutations are arising uniformly in time, and the time at which a mutation is observed is random, the SFS is proportional to the average time an allele is expected to spend in a given frequency window.

In this section, we show that the low and high frequency ends of the site frequency spectrum of unlinked loci can be obtained from a simple heuristic argument that emphasizes this connection between allele frequency trajectories and the site frequency spectrum. These calculations are not intended to be exact, but they provide intuition for the origins of key features of the site frequency spectrum that we will return to more formally below.

Consider the simplest case of unlinked purely neutral loci. Neutral mutations will arise in the population at rate *NU*_*n*_. In the absence of selection, the trajectories of these mutations are governed by genetic drift. At steady state, the number of mutations we expect to see at frequency *f* is simply proportional to the number of mutations that reach that frequency and the typical time each of these mutations spend at that frequency before fixing or going extinct. In the absence of selection, a new mutation that arises at initial frequency 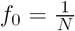 will reach frequency *f* before going extinct with probability 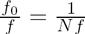. Standard branching process calculations (Fisher, 2007) show that, given that it reaches frequency f, the mutation will spend about *Nf* generations around that frequency (defined as log(*f*) not changing by more than 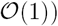, provided that *f* is small (*f* ≪ 1).

By combining these results, we can calculate the expected site frequency spectrum for small *f*. The rate at which new mutations reach frequency *f* is 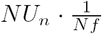. Those that do will remain around *f* (in the sense defined above) for about *Nf* generations. Thus the total number of neutral mutations within *df* of frequency *f* is 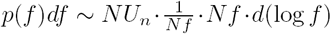. In other words, we have

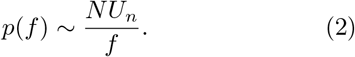

This argument is valid when *f* is rare, but will start to break down at intermediate frequencies. However, because the wild type is rare when the mutant approaches fixation, an analogous argument can be used to describe the site frequency spectrum at high frequencies. The mutant trajectory still reaches frequency *f* with probability 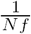. It will then spend roughly 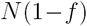 generations around this frequency (i.e. within 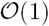 of log(1 — *f*)). This gives 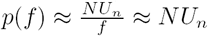 in the high frequency end of the spectrum. This simple forward-time heuristic argument reproduces a well-known result of coalescent theory (Wakeley, 2009) and agrees with the more formal calculation of sojourn times in the Wright-Fisher process (Ewens, 1963).

We can use a similar argument to calculate the frequency spectrum of strongly selected deleterious mutations with fitness effect -*s* (with *Ns* ≫ 1) that occur at a locus that is unlinked to any other selected locus. Provided that the deleterious mutation is rare (below the “drift barrier” frequency, 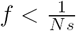), its trajectory is dominated by drift. Thus for 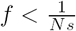, the mutation trajectory will be the same as for a neutral mutation, and the frequency spectrum will therefore be neutral. In contrast, at frequencies larger than 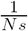 selection is stronger than drift, which prevents the mutation from exceeding this frequency. Combining these two expressions, we find that the frequency spectrum of an unlinked deleterious mutation is to a rough approximation given by

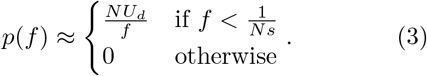

For completeness, we also show how a similar argument can be used to obtain the frequency spectrum of beneficial mutations. Though it is not immediately obvious that this is relevant to background selection, we will later see how similar trajectories emerge in the case of strong purifying selection. Just like deleterious alleles, strongly beneficial alleles with fitness effect *s* (with *Ns* ≫ 1) will not feel the effects of selection as long as they do not exceed the drift barrier 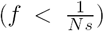. Their trajectory and frequency spectrum will therefore be neutral below the drift barrier. As a result, only a small fraction *s* of beneficial mutations will reach frequency 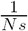. However, those that do will be destined to fix, since at frequencies larger than 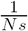, selection dominates over drift. Above this threshold, selection will cause the frequency of the mutation to grow logistically at rate 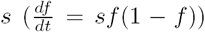, spending 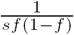 generations near frequency *f*. This is valid as long as 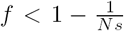, at which point the effects of drift become dominant due to the wild type being rare, and the trajectory of the mutant is once again the same as the trajectory of a neutral mutation. Combining these expressions, we obtain a rough approximation for the frequency spectrum of an unlinked beneficial mutation

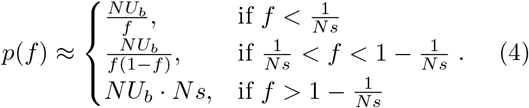

## BACKGROUND SELECTION MODEL

We now turn to the analysis of background selection. We focus on the simplest possible model of purifying selection at a perfectly linked genetic locus in a population of *N* individuals. We assume neutral mutations occur at a per-locus per-generation rate *U*_*n*_, and deleterious mutations occur at rate *U*_*d*_ (*U*_*d*_ ≪ 1). Throughout the bulk of the analysis, we will assume that all deleterious mutations reduce the (log) fitness of the individual by the same amount *s* (*s* ≪ 1), though we analyze the effects of relaxing this assumption in a later section below. We neglect epistasis throughout, so that the fitness of an individual with *k* deleterious mutations at this locus is *–ks.* For simplicity we consider haploid individuals, but our analysis also applies to diploids in the case of semidominance (*h* = 1/2).

Our model is equivalent to the non-epistatic case of the model formulated by Kimura and Maruyama (1966) and Haigh (1978) and to the 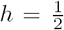 case of the model considered by Charlesworth et al. (1993) and Hudson and Kaplan (1994), and later studied by many other authors (Gordo et al., 2002; Nicolaisen and Desai, 2012; Seger et al., 2010; Walczak et al., 2011). However, instead of modeling the genealogies of a sample of individuals from the population backwards in time, we offer a forward-time analysis of this model, in which we analyze the full frequency trajectory of alleles.

We define *λ* = *U*_*d*_*/s*, and we assume that selection is sufficiently strong that alleles carrying deleterious mutations cannot fix in the population (*Nse*^−*λ*^ ≫ 1). The opposite case, in which deleterious mutations are weak enough to routinely fix (*Nse^−λ^ ≲* 1), leads to the rapid accumulation of deleterious mutations via Muller’s ratchet; we and others have studied mutational trajectories and genealogies in this limit in earlier work (Good and Desai, 2013; Good et al., 2014; Neher and Hallatschek, 2013). In the Discussion we comment on the connection between these earlier weak-selection results and the strong-selection case we study here.

Since we assume that all mutations have the same effect on fitness, the population can be partitioned into discrete fitness classes according to the number of deleterious mutations each individual carries at the locus. When the fitness effect of each mutation is sufficiently strong, the population assumes a steady-state fitness distribution in which the expected fraction of individuals with *k* deleterious mutations, *h*_*k*_, follows a Poisson distribution with mean 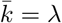 (Haigh, 1978; Kimura and Maruyama, 1966),

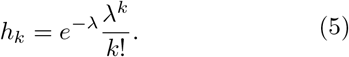

A new allele in such a population will arise on a background with *k* existing mutations with probability *h*_*k*_.

From the form of *h*_*k*_, we see that depending on the value of λ, the population can be in one of two regimes. In the first regime, the rate at which mutations are generated is smaller than the rate at which selection can purge them (λ ≪ 1). In this case, the majority of individuals in the populations carry no deleterious mutations (*h*_0_ ≈ 1), with only a small proportion, 1 – *h*_0_ ≈ λ, of backgrounds in the population carrying some deleterious variants. To leading order in λ, all new neutral mutations will arise in a mutation-free background and will remain at the same fitness as the founding genotype. Their trajectories are thus the same as the trajectories of mutants at unlinked genetic loci of the same fitness as the founding genotype (see Appendix D for details). This means that the full site frequency spectrum can be calculated by summing the contributions of unlinked site frequency spectra that we calculated above. The neutral and deleterious site frequency spectra are, to leading order in λ, given by Equations 2 and 3, respectively (for details see Appendix H). Thus, background selection has a negligible impact on mutational trajectories and diversity when λ ≪ 1.

In the opposite regime where λ ≫ 1, mutations are generated faster than selection can purge them and there will be substantial fitness variation at the locus. Consider a new allele (i.e. a new mutation at some site within the locus) that arises in this population. A short time after arising, individuals that carry this allele will accumulate newer deleterious mutations, which will lead the allele to spread through the fitness distribution. The fundamental difficulty in calculating the frequency trajectory of this allele, *f*(*t*), stems from the fact that a short time after arising, individuals that carry the allele will have accumulated different numbers of newer deleterious mutations. The total strength of selection against the allele depends on the average number of deleterious mutations that the individuals that carry the allele have. This will change over time in a complicated stochastic way, as the lineage purges old deleterious mutations, accumulates new ones, and changes in frequency due to drift and selection. To calculate the distribution of allele frequency trajectories in this regime, we will need to model these changes in the fitness distribution of individuals carrying the allele. Although we will formally be treating *λ* as a large parameter, in practice our results will also adequately describe allele frequency trajectories in the cases of moderate *λ* (i.e. *λ* ≳ 3).

To make progress, we classify individuals carrying this allele (the “labeled lineage”) according to the number of deleterious mutants they have at the locus. We denote the total frequency of the labelled individuals that have *i* deleterious mutations as *f*_*i*_(*t*), so that the total frequency of the lineage, *f*(*t*), is given by

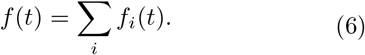

The time evolution of the allele frequency in a Wright-Fisher process is commonly described by a diffusion equation for the probability density of the allele frequency (Ewens, 2004). Instead, for our purposes it will be more convenient to consider the equivalent Langevin equation (Van Kampen, 2007),

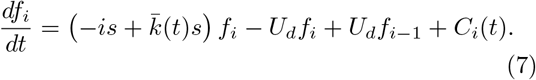

Here, 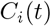 is a noise term with a complicated correlation structure necessary to keep the total size of the population fixed (see Good and Desai (2013) for details) and 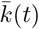 is the mean number of mutations per individual in the entire population at time *t.* In the strong selection limit that we are interested in here (*Nse*^−*λ*^ ≫ 1), fluctuations in the mean of the fitness distribution of the population are small and 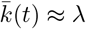 (Neher and Shraiman, 2012).

## KEY FEATURES OF LINEAGE TRAJECTORIES

Before turning to a detailed analysis of Eq. 7, it is helpful to consider some of the key features of lineage trajectories that we will model more formally below. To begin, imagine a lineage founded by a neutral mutation in an individual with *k* deleterious mutations. Let the lineage be composed of *n*_*k*_(0) individuals at some time *t* = 0 shortly after arising, all of which carry *k* deleterious mutations (see blue inset in Figure 2A). At this time, the relative fitness of this lineage is simply 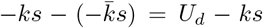. Thus, lineages founded in classes with *k* > *λ* will tend to decline in size. In contrast, the more interesting case arises if *k* < *λ*, since these lineages will tend to increase in size.

**FIG. 2.**
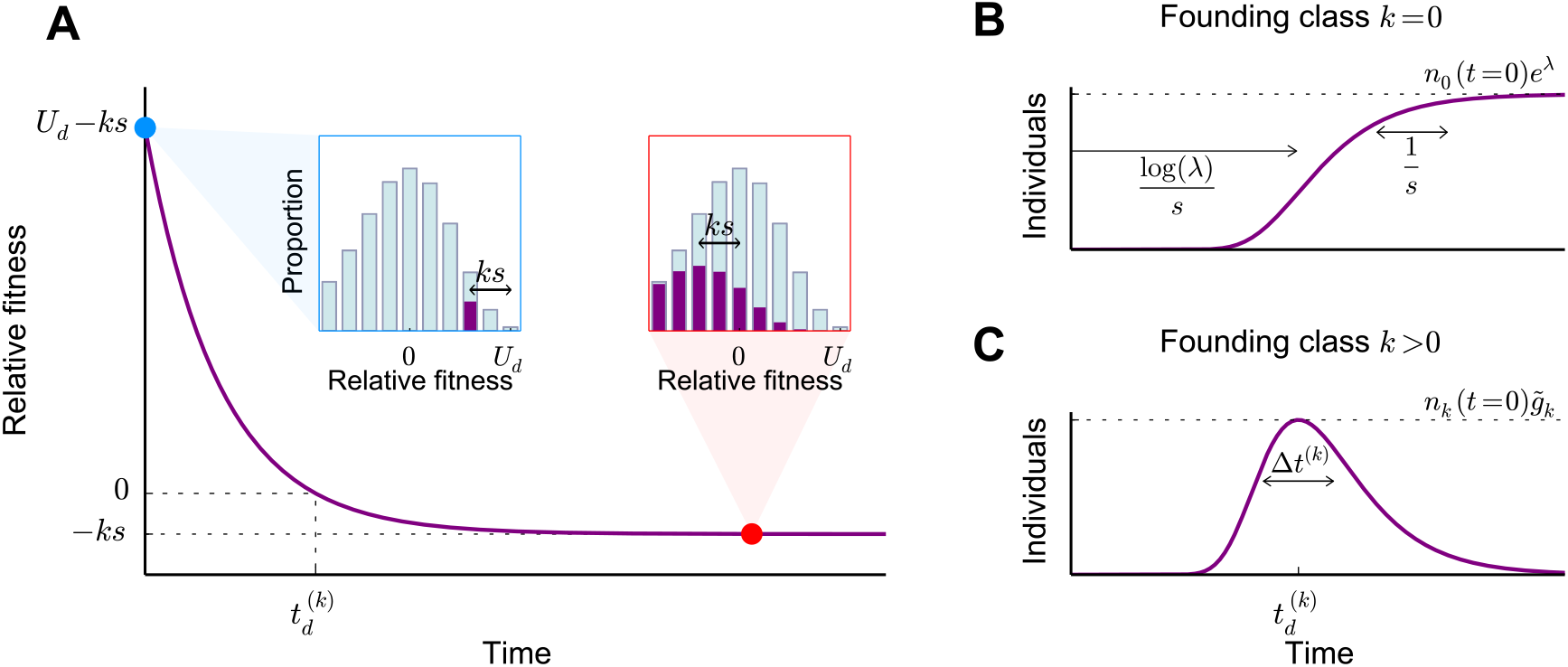
(A) The average fitness of a lineage composed of individuals carrying *k* deleterious mutations at time *t* = 0 (blue dot and blue inset). As the descendants of these individuals accumulate further deleterious mutations, the fitness of the lineage declines until the individuals accumulate an average of *λ* deleterious mutations (red dot) and reach their own mutation-selection balance, which is a steady-state Poisson profile with mean *λ* that has been shifted by the initial deleterious load, *−ks*, (red inset). (B) In the absence of genetic drift, the lineage will increase at a rate proportional to its relative fitness. Lineages arising in the class with *k* = 0 deleterious mutations reach mutation-selection balance about 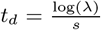 after arising, after which the size of the lineage asymptotes to *n*_0_ (*t* = 0)*e*^*λ*^. The fitness of these lineages changes on a shorter timescale, 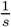. (C) In contrast, lineages arising in classes with *k* > 0 deleterious mutations peak in size after a time 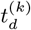, when their average relative fitness is zero, after which they decline exponentially at rate *ks.*

However, though the overall number of individuals that carry the allele will tend to increase when *k* < *λ*, the part of the lineage in the founding class *k* (the 'founding genotype') will tend to decline in size, because it loses individuals through new deleterious mutations (at per-individual rate – *U*_*d*_). As a result, the founding genotype feels an effective selection pressure of *U*_*d*_ – *ks* – *U*_*d*_ = *–ks*, which is negative for all *k* > 0 and 0 for *k* = 0. This means that the lineage will increase in frequency not through an increase in size of the founding genotype, but rather through the appearance of a large number of deleterious descendants in classes of lower fitness. The lineage must therefore decline in fitness as it increases in size.

In the absence of genetic drift, we can calculate how the size and fitness of the lineage change in time by dropping the stochastic terms in Eq. 7 (subject to the initial condition *n*_*k*_(*t* = 0) = *n*_*k*_(0) and *n*_*k+i*_(*t* = 0) = 0 for all *i* ≠ 0). These deterministic dynamics of the lineage have been analyzed previously by Etheridge et al. (2007), who showed that the number of additional mutations that an individual in the lineage carries at some later time *t* is Poisson-distributed with mean *λ*(l – *e*^*–st*^). Thus the average number of additional deleterious mutations eventually approaches *λ* after 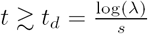 generations. At this point, the lineage has reached its own mutation-selection balance: the fitness distribution of the lineage has the same shape as the distribution of the population (i.e. *n*_*i+k*_(*t*) = *h*_*i*_ · *n*_*k*_(*t*)), but is shifted by – *ks* compared to the distribution of the population (see red inset in Figure 2A).

The average relative fitness *x*(*t*) of individuals in the lineage (Fig. 2A) is therefore equal to

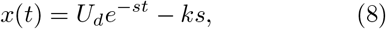

and the total number of individuals in the lineage is simply 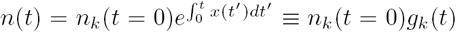, where we have defined

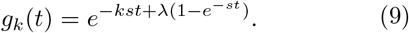

Thus, we can see from Equations 8 and 9 that lineages founded in the 0-class will on average steadily increase in size at a declining rate until they asymptote at a total size equal to *n*_*k*_(*t* = 0)*e*^*λ*^ roughly 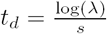 generations later (see Figure 2B). In contrast, lineages founded in the *k*-class will increase in size for only

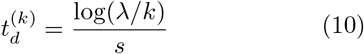

generations, when they peak at a size of 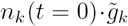 individuals (see Figure 2C), where we have defined

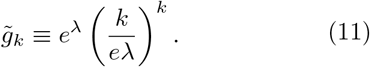

The lineages remain near this peak size for about

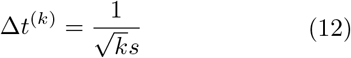

generations (Fig. 2C). At longer times, they exponentially decline at rate –*ks* (Fig. 2C).

These simple deterministic calculations capture the average behavior of an allele and show that all alleles founded in classes with *k* > 0 are likely to be extinct on timescales much longer than 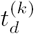, whereas sufficiently large lineages founded in the 0-class should simply reflect the frequency in the founding class about *t*_*d*_ generations earlier, 〈*f*(*t*)) ≈ *e*^*λ*^*f*_0_(*t*–*t*_*d*_). This is the forward-time analogue of the intuition presented by Charlesworth et al. (1993).

Of course, this deterministic solution neglects the effects of genetic drift, which will be crucial, particularly because drift in each class propagates to affect the frequency of the lineage in all lower-fitness classes (for a more detailed heuristic describing why drift can never be ignored, see Appendix B). Although these effects are complex, there is a hierarchy in the fluctuation terms which we can exploit to gain some intuition. From the deterministic solution above, we can see that a fluctuation of size *δf*_*i*_ in class *i* will on average eventually cause a change in the total size of the lineage proportional to 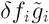 after a time delay 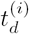. Thus, the fluctuations that have the largest effect on the total size of the lineage are those that occur in the class of highest fitness (i.e. the founding class *k*). These fluctuations will turn out to be the most important to describing the frequency trajectory of the entire allele, though fluctuations in classes of lower fitness will still matter in lineages of small enough size.

One could imagine that this result means that fluctuations in the total size of the lineage simply mirror the fluctuations in the founding class, amplified by a factor 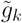 and after a time delay 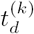. If fluctuations in the founding class are sufficiently slow, this is indeed the case. However, for fluctuations that occur on shorter timescales, this is not true. Consider for example the case where a neutral mutation is founded in the mutation-free (*k* = 0) class. Imagine that the frequency of the allele in the founding class changes by a small amount from *f*_0_ to *f*_0_ + *δf*_0_ as a result of genetic drift (shown in the first panel of Figure 3). Based on the deterministic solution, this fluctuation will lead to a proportional change in the frequency of the portion of the lineage in the 1-class, and this change will take place over ~ 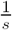 generations (see Appendix A for details). During this time, the change in the 1-class begins to lead to a shift in the frequency in the 2-class, which will mirror the change in the 0-class a further 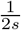 generations later (see Fig. 3). This change will then propagate in turn to lower classes, and ultimately results in a proportional change in the total allele frequency a total of 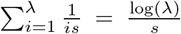 generations later (see Fig. 3).

**FIG. 3.**
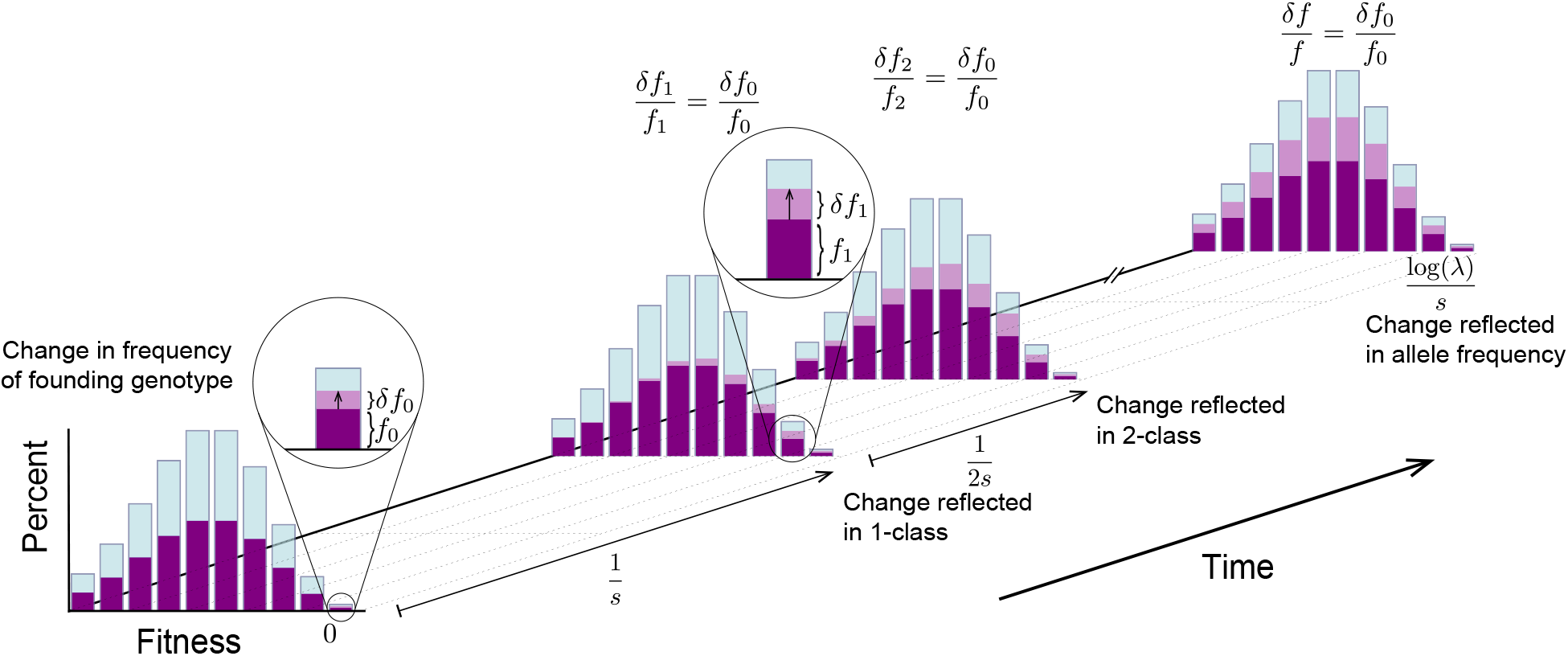
A schematic showing how a change in the frequency of the lineage in the mutation-free class propagates to affect the frequency in all classes of lower fitness. At time *t* = 0, the lineage is in mutation-selection balance at total frequency *f*, when the frequency of the portion of the lineage in the zero class changes suddenly from *f*_0_ to *f*_0_ + δ*f*_0_. This change is felt in the 1-class 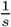 generations later, and propagates to the 2-class yet another 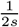 generations later. The lineage reaches a new equilibrium about 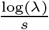 generations later, when the total allele frequency is proportional to (*f*_0_, + *δf*_0_)*e*^*λ*^.

Now consider what happens if there is another change in the frequency in the founding class. If this change occurs within the initial 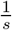 generations, it will influence the 1-class simultaneously with the first fluctuation, and thus the effect of these two fluctuations on the overall lineage frequency will be “smoothed” out. In contrast, if the changes are separated by more than 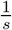 generations, they will propagate sequentially through the fitness distribution, and are ultimately mirrored in the total allele frequency. Similar arguments apply to lineages founded in other fitness classes, though the relevant timescales and scale of amplification are different.

Together, these arguments suggest that fluctuations in the founding class will have the largest impact on overall fluctuations in the lineage frequency, and these overall fluctuations will represent an amplified but smoothed-out mirror of the fluctuations in the founding class. This smoothing will be crucial: the size of the lineage in the founding class will typically fluctuate neutrally, but the smoothed-out and amplified versions will have non-neutral statistics. As we will see below, this smoothing ultimately leads to distortions in the site frequency spectrum at low frequencies 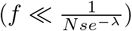.

## ANALYSIS

Formally, we analyze all of the effects described above by computing the distribution of the frequency trajectories *f*(*t*) of the allele, *p*(*f*, *t*), from Eq. 7 for an allele arising in class *k.* This process is complicated by the correlation structure in the *C*_*i*_(*t*) terms required to keep the population size constant. These correlations are important once the lineage reaches a high frequency, and in the presence of strong selection they result in a complicated hierarchy of the moments of f, which do not close (Good and Desai, 2013; Higgs and Woodcock, 1995). However, we can simplify the problem by considering low, high, and intermediate-frequency lineages separately. First, at sufficiently low frequencies (*f* ≪ 1), the *C*_*i*_(*t*) in Eq. 7 reduce to simple uncorrelated white noise. At these low frequencies, Eq. 7 thus simplifies to

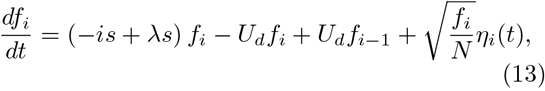

where the noise terms have 〈*η*_*i*_(*t*)〉 = 0 and covariances 〈*η*_*i*_(*t*)*η*_*j*_(*t*′)〉 = *δ_ij_δ*(*t* — *t′*) and should be interpreted in the Itô sense. At very high frequencies (1 — *f* ≪ 1), a similar simplification arises. In this case, the wild-type lineage is at low frequency, and we can model the wild-type frequency using an analogous coupled branching process with uncorrelated white noise terms. Finally, at intermediate frequencies, we cannot simplify the noise terms in this way. Fortunately, for the case of strong selection we consider here, we will show that for 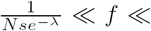 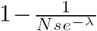, lineage trajectories have neutral statistics on relevant timescales. As we will see below, these low, intermediate, and high frequency solutions can then be asymptotically matched, giving us allele frequency trajectories and site frequency spectra at all frequencies.

In the next several subsections, we focus on the analysis of the distribution of trajectories at low and high frequencies (*f* ≪ 1 or 1 – *f* ≪ 1), where Eq. 13 is valid. We then return in a later subsection to the analysis of trajectories at intermediate frequencies.

### The dynamics of the lineage within each fitness class

To obtain the distribution of trajectories of the allele *p*(*f*,*t*) at low frequencies (*f* ≪ 1) from Eq. 13, we will first compute the generating function of *f*(*t*). This generating function is defined as

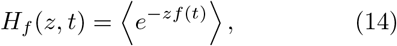

where angle brackets denote the expectation over the probability distribution of the frequency trajectory *f*(*t*). *H*_*f*_(*z*,*t*) is simply the Laplace transform of the probability distribution of *f*(*t*), and it therefore contains all of the relevant information about the probability distribution of *f*(*t*).

As we have already anticipated from our discussion above, the time evolution of *f*(*t*) depends on the distribution of the lineage among different fitness classes. In order to understand how this distribution changes under the influence of drift, mutation and selection in these classes, we can consider the joint generating function for the *f*_*i*_(*t*),

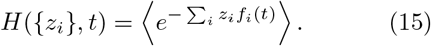

The generating function for the total allele frequency *H*_*f*_(*z*, *t*) can then be obtained from this joint generating function by setting *z*_*i*_ = *z.* We will use this relationship between the two generating functions to evaluate the importance of drift, mutation and selection within each of the fitness classes on the total allele frequency.

By taking a time derivative of Eq. 15 and substituting the time-derivatives 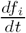 from Eq. 13 (where the stochastic terms should be interpreted in the Itô sense), we can obtain a partial differential equation describing the evolution of the joint generating function,

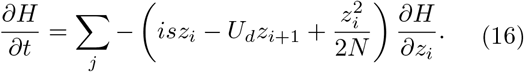

We see from Equation 16 that the joint generating function is constant along the characteristics *z*_*i*_(*t – t′*) defined by

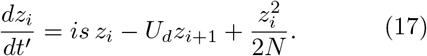

Thus, the joint generating function can be obtained by integrating along the characteristic backwards in time from *t′* = 0 to *t′* = *t*, subject to the boundary condition *z*_*i*_(*t*) = *z.* Note that the linear terms in the characteristic equations arise from selection and mutation into and out of the *i* class, and that the nonlinear term arises from drift in class *i.*

In Appendix E, we show that when considering the distribution of trajectories *p*(*f*, *t*) at frequencies 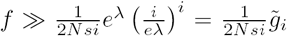 the nonlinear terms in Eq. 17 are of negligible magnitude uniformly in time in all classes containing *i* or more deleterious mutations per individual as long as *i* ≪ *λ*, *Nse*^−*λ*^ ≫ 1 and *λ* ≫ 1. Here, 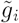 represents the peak of the expected number of individuals in a lineage founded by a single individual in class *i* (see Eq. 11 and Fig. 2C). Thus, when 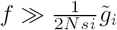, the effect of genetic drift is negligible in classes with *i* or more deleterious mutations. Conversely, when 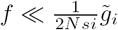, genetic drift in the class with *i* deleterious mutations does affect the overall allele frequency.

Since drift is negligible in classes with *i* or more mutations, total allele frequencies of 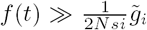 require that 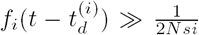. This threshold is reminiscent of the 'drift barrier', but its origin for classes below the founding class (*i* > *k*) is more subtle. We offer an intuitive explanation for this threshold in Appendix B. Thus, drift in class *i* has an important impact on the overall frequency trajectory as long as 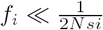. However, once *f*_*i*_ exceeds 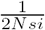, the effect of genetic drift in that class, as well as all classes below *i*, becomes negligible, because the frequencies of the parts of the lineage in all classes below *i* are then also guaranteed to exceed the corresponding thresholds. Note that because the frequency of the founding genotype *f*_*k*_ is exponentially unlikely to substantially exceed 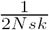, the overall allele frequency *f* is exponentially unlikely to substantially exceed 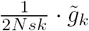.

In summary, by analyzing the generating function for the components of the lineage in different fitness classes, we have found that there is a clear separation between high fitness classes in which mutation and drift are the primary forces, and classes of lower relative fitness in which mutation and selection dominate. The boundary between the stochastic and deterministic classes can be determined from the total allele frequency, allowing us to reduce a complicated problem involving a large number of coupled stochastic terms to what we will see is a small number of stochastic terms feeding an otherwise deterministic population.

### Statistics of trajectories with 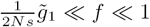

At this point, we are in a position to calculate a piecewise form for the generating function *H*_*f*_(*z*, *t*) valid near any frequency *f*. For example, consider the allele frequency trajectory in the vicinity of some frequency 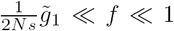. As we have explained above, at these frequencies contributions from individuals arising in class *k* ≥ 1 are exponentially small, since they would require the frequency of the lineage in that class to substantially exceed 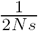, which happens only exponentially rarely. Thus, in this frequency range we will only see mutations arising in the mutation-free class (*k* = 0). In addition to this, we have shown that at these frequencies genetic drift can be neglected in all classes but the 0-class. To obtain the generating function at these frequencies, we can therefore integrate the characteristic equations by dropping the nonlinear terms in Eq. 17 for all *i* > 0 (see Appendix E for details). This yields the generating function for the frequency of the labelled lineage,

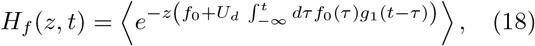

where the average is taken over all possible realizations of the trajectory in the founding class *f*_0_(*t*).

As before, *g*_1_(*t –* τ) represents the expected number of individuals descended from an individual present in class in the 1-class *t – τ* generations earlier (see Eq. 9). Thus, the two terms in the exponent in Eq. 18 represent the frequency of the lineage in the founding class *f*_0_ and the total frequency of the deleterious descendants of that lineage. The latter are seeded into the 1-class at rate *NU_d_ f*_0_(*τ*), and each of these deleterious descendants founds a lineage that *t –* τ generations later contains *g*_1_(*t* – τ) individuals, so that the total frequency of the allele is simply

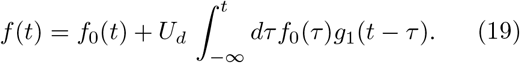

Thus, we have obtained a simple expression for the frequency of the entire allele in which all of the stochastic effects have been reduced to a single stochastic component, *f*_0_(*t*). Furthermore, the stochastic dynamics of *f*_0_(*t*) are those of a simple, unlinked neutral mutation (see schematic of such a trajectory in Figure 4B). Note however that the statistics of the fluctuations in *f*(*t*) are *not* necessarily the same as the statistics of the trajectory in the founding class, because the trajectory *f*_0_(*t*) is correlated in time (see Fig. 4B), and these correlations, when integrated in time, will lead to the total frequency trajectory *f*(*t*) of the lineage having different stochastic properties (see Figure 4A).

**FIG. 4.**
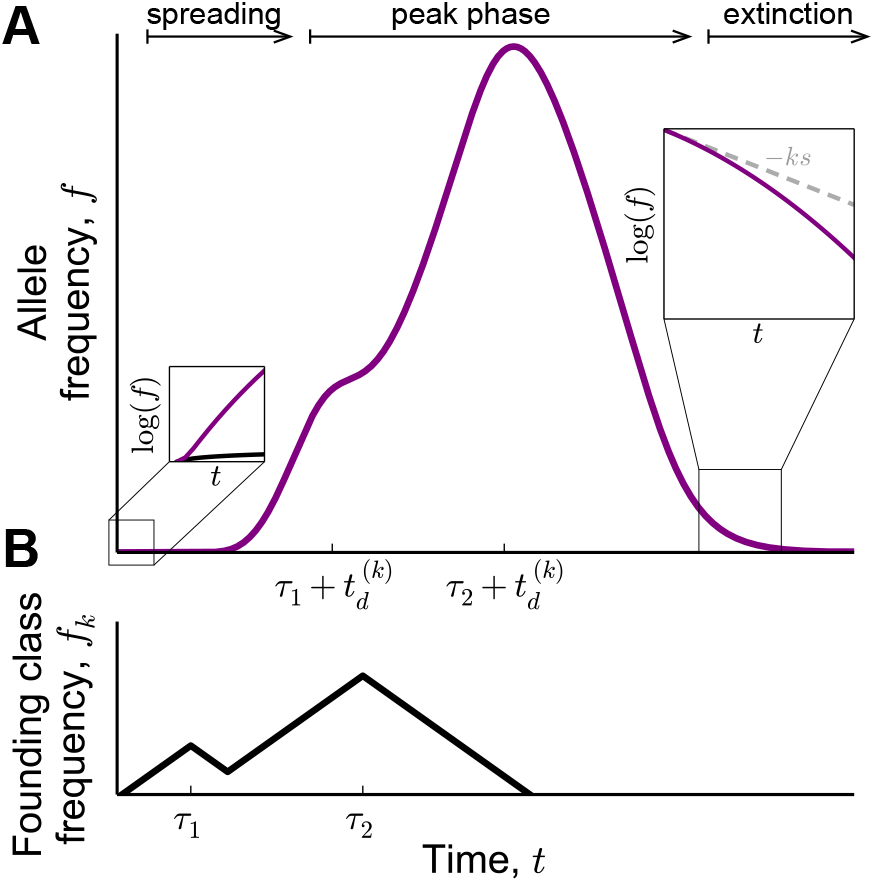
Schematic of (A) the trajectory of the total allele frequency, and (B) the trajectory of the frequency of the portion of the allele that remains in the founding class. Soon after arising in the founding class, the allele frequency rapidly increases in the spreading phase of the trajectory. Early in this phase, the total allele frequency (purple) becomes much larger than the frequency of the founding genotype (black; left inset, panel A). In the peak phase of the trajectory, the total allele frequency trajectory represents a smoothed out and amplified version of the trajectory in the founding class. In the extinction phase of the trajectory, the allele frequency declines at an increasing rate (right inset, panel A).

From Eq. 19, we can see that the frequency trajectory of the allele still has the same qualitative features as those we have seen in the deterministic behavior of mutations. Shortly after being founded, the lineage will become dominated by the deleterious descendants of the founding class, which are captured by the second term in Eq. 19 (see left inset in Fig. 4A). At early times 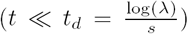, the total allele frequency must rapidly grow, as the lineage spreads through the fitness distribution and approaches mutation-selection balance (see Figure 4A). About *t*_*d*_ generations after founding, the peak phase of the trajectory begins (see Fig. 4A). During this phase, the average fitness of the lineage is approximately 0 and the allele traces out a smoothed-out and amplified version of the trajectory in the founding class (Fig. 4B). Finally, *t*_*d*_ generations after the descendants of the last individuals present in the founding class have peaked, the average fitness of the lineage will fall significantly below 0, and the extinction phase of the trajectory begins.

As we show in Appendix H, the peak phase of the trajectory is the most important for understanding the site frequency spectrum. This is also the phase during which the trajectory of the mutation spends the longest time near a given frequency. In contrast, the spreading phase (see Fig. 4A) has a negligible effect on the site frequency spectrum: by this we mean that the site frequency spectrum at a given frequency will always be dominated by the peak phase of trajectories that peak around that frequency, and not influenced by the spreading phase of trajectories that peak at much higher frequencies. We will therefore not consider the spreading phase in the main text, but discuss it in Appendix H. The extinction phase of the trajectory can also be neglected for a similar reason (Appendix H), except when considering the very highest frequencies, 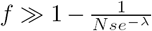. At these frequencies, the wild type frequency is small and the mutant is in the process of fixation. To analyze the allele frequency trajectory at these frequencies, we model the wild type using the coupled branching process in Eq. 13, and hence describe these trajectories by the extinction phase of the wild type.

To calculate the distribution of *f*(*t*) in the peak phase, we need to calculate the distribution of this time integral of *f*_0_(*t*) in Eq. 19. We can simplify this integral by observing that *g*_1_(*t*) is highly peaked in time between 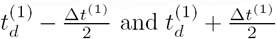, where 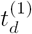 and Δ*t*^(1)^ are given by Equations 10 and 12 and annotated in Figure 2C. In other words, starting at times around 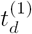 generations after the lineage reaches a substantial frequency in the founding class, the labelled lineage is dominated by the deleterious descendants of individuals extant in the founding class between 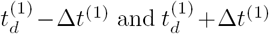 generations earlier, with individuals extant in the founding class at other times having exponentially smaller contributions (see Appendix E for details). Thus, the total size of the lineage will be proportional not to the frequency 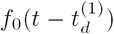 in the founding class 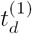 generations earlier, but to the total time-integrated frequency within some window of width ~ Δ*t*^(1)^ centered around that time. We call this quantity the “weight” and denote it by *W*_*Δt*^(1)^_, where

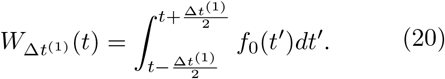

The total allele frequency in the peak phase is therefore equal to

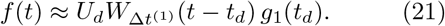

Thus, to calculate the distribution of the allele trajectory, we only need to calculate the distribution of the weight in the founding class over a window of specified width, Δ*t*^(1)^. It is informative to consider the time-integrated form of the distribution of this weight, 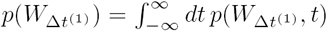, since this form is also directly relevant to the site frequency spectrum (for a discussion of the time-dependent distribution *W*_Δ*t*^(1)^_(*t*), see Appendix F). In Appendix F we show that *p*(*W*_*Δt*^(1)^_) is given by

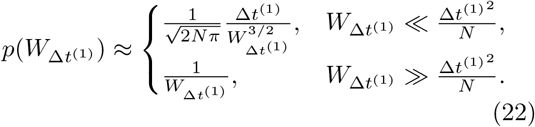

This distribution has a form that can be simply understood in terms of the trajectory in the founding class. Since genetic drift takes order *N f*_0_ generations to change *f*_0_ substantially, drift will not change *f*_0_ significantly within Δ*t*^(1)^ generations when the frequency in the founding class exceeds 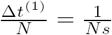. As a result, the weight, *W*_Δ*t*(1)_, will be approximately equal to 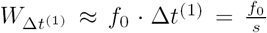. Therefore, at these large frequencies, the weight simply traces the feeding class frequency, and the two quantities have the same distributions. At lower frequencies, 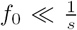, the founding genotype will typically have arisen and gone extinct in a time of order *N f*_0,max_ generations (where *f*_0,max_ the maximal frequency the lineage reaches over the course of its lifetime). By assumption, this time is much shorter than 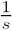. Thus, the weight in a window of width 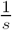 that contains this trajectory is simply 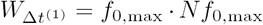. This large a trajectory is obtained with probability 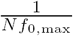, from which it fol-lows (by a change of variable) that the distribution of weights in the founding class scales as 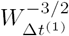.

As we anticipated in our discussion of the propagation of fluctuations of the founding genotype through the fitness distribution (Fig. 3), we have found that the trajectory of the allele in the peak phase looks like a smoothed out, time-delayed and amplified version of the trajectory in the founding class (Fig. 4). At sufficiently high frequencies, 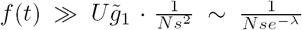, the timescale of the smoothing is shorter than the typical timescale of the fluctuations in the founding class. At these frequencies the statistics of the fluctuations of the allele simply mirror the statistics of the fluctuations in the founding class, with a time delay equal to 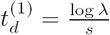.

At lower frequencies, 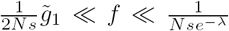, the timescale of smoothing is much longer than the typical lifetime of the founding genotype. As a result, the deleterious descendants of the entire original genotype rise and fall simultaneously and fluctuations in the founding class are not reproduced in detail. Instead, the peak phase of the allele frequency trajectory consists of a single peak with size proportional to the total lifetime weight of the founding genotype, 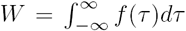. As we calculated above, the distribution of these peak sizes falls off more rapidly than neutrally. This gives us a complete description of the statistics of the peaks of allele frequency trajectories in the frequency range 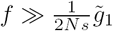.

### Statistics of trajectories with 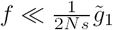

So far, we have only considered trajectories of lineages that reach a maximal allele frequency larger than 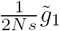, all of which must have arisen in the mutation-free class. At lower frequencies, 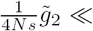 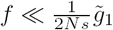, the effects of genetic drift in class *i* = 1 must also considered, but the behavior in classes with *i* ≥ 2 is deterministic. In this case, by repeating our earlier procedure, we obtain a slightly different form for the generating function *H*_*f*_(*z*, *t*),

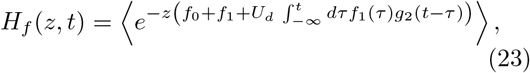

so that the total allele frequency is

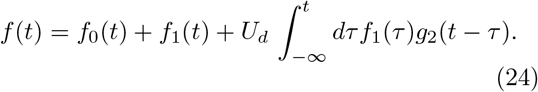

The total allele frequency is once again dominated by the last term, which represents the bulk of the deleterious descendants. Thus, by an analogous argument, the peak size of the lineage is proportional to the weight in the 1-class in a window of width 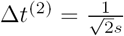,

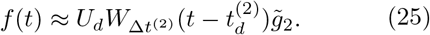

There are two types of trajectories that can reach these frequencies: trajectories that arise in the 1-class and reach a sufficiently large frequency in their founding class (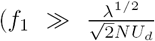, see Appendix G), and trajectories that arise in the 0-class and reach a smaller frequency in their founding class (*f*_0_ ≪ 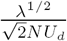), but still leave behind enough deleterious descendants in the 1-class that the overall frequency in that class exceeds 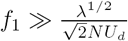. By the argument that we outlined before, this ensures that genetic drift will negligible in classes of lower fitness (i.e. for *i ≥* 2), and is guaranteed to happen if 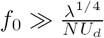 (see Appendix G).

The trajectories of the former type are simple to understand, since in this case, the trajectory *f*_1_(*t*) is that of a simple, unlinked deleterious locus with fitness *–s* (and *f*_0_(*t*) = 0 at all times). By repeating the same procedure as above, we find that the time-integrated distribution of the weights in the 1-class is

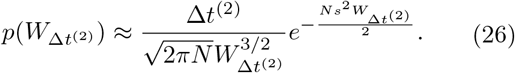

Note that since the trajectory of a mutation in the founding 1-class is longer than ~ 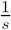 generations only exponentially rarely, a window of length Δ*t*^(2)^ nearly always contains the entire founding class trajectory (see Appendix F). This is reflected in the form of the weight distribution in Equation 26, which falls as 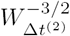 with an exponential cutoff at 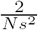. Thus, the frequency trajectory of an allele that arises in the 1-class will not mirror the fluctuations in the founding genotype. Instead, the peak phase of the allele frequency trajectory will nearly always consist of a single peak, just as we have seen in the case of alleles peaking at frequencies 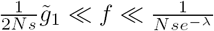.

We now return to the other type of trajectory that can peak in this range: alleles arising in the 0-class, but reaching a small enough frequency that the effects of genetic drift in the 1-class cannot be ignored 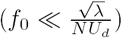. Because the trajectory of these alleles in the 1-class represents the combined trajectory of multiple clonal sub-lineages each founded by a mutational event in the 0-class, the distribution of weights in the 1-class will be different 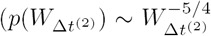, see Appendix G), which leads to a different distribution of overall allele frequencies *f*. However, as we show in Appendix H, these trajectories have a negligible impact on the site frequency spectrum: because the overall number of mutations arising in class 1 is substantially larger than the overall number of mutations arising in class 0, trajectories that arise in class 0 and peak in the same frequency range as mutations originating in class 1 are less frequent by a large factor (*λ*^3/4^, see Appendix G),.

Similarly, at even lower frequencies in the range 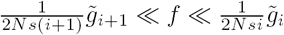, we will see the peaks of trajectories arising on backgrounds with *i* or fewer deleterious mutations. These trajectories all have a single peak of width equal to 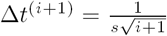. The maximal peak sizes are, once again, proportional to the total weight in the *i*-class, which will be distributed according to a different power law depending on the difference in the number of deleterious mutations Δ = *k – i* between the founding class *k* and the *i*-class (see Appendix G for details), with the most numerous ones being those that arise in the *i*-class (*k* = *i*) (Appendix H).

At the very lowest frequencies, 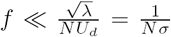, the site frequency spectrum is dominated by the trajectories of lineages that arise in a class that is within a standard deviation *σ* of the mean of the fitness distribution (i.e. lineages with 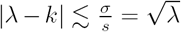. Unlike the trajectories of lineages that arise in classes of higher fitness that we discussed above, allele frequency trajectories of lineages arising within a standard deviation of the mean are typically dominated by drift throughout their lifetime (see Appendix E). This is because the timescale on which these lineages remain above the mean of the fitness distribution (which is limited by *t*_*d*_(*k*)) is shorter than the timescale that it takes them to drift to a frequency large enough for the effect of selection to be felt 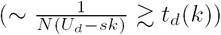. Lineages arising in these classes do not reach frequencies substantially larger than 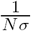, and have largely neutral trajectories at frequencies that remain below this threshold.

### The mirrored fluctuations of the allele at intermediate frequencies, 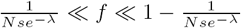

We have seen that the effects of genetic drift in multiple fitness classes may be important when *f* ≪ 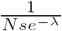, but that at frequencies larger than 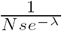, genetic drift in all classes apart from the 0-class can be neglected and the trajectory of the allele mirrors the fluctuations in the 0-class that occur on timescales longer than 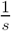 generations.

Once an allele reaches a more substantial frequency, the coupled branching process in Eq. 13 no longer adequately describes its allele frequency trajectory. However, as long as both the mutant and wild-type frequencies remain larger than 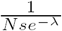, the effects of drift in all classes apart from the 0-class will remain negligible and we can use a coarse-grained model to describe the allele frequency trajectories on timescales longer than 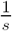.

On these timescales, we can approximate the total allele frequency of both the mutant and the wild type frequencies as 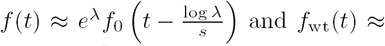 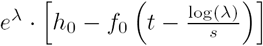, which yields a model for the total allele frequency of the mutant

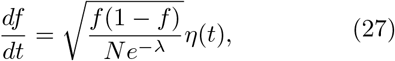

where *η*(*t*) is an effective noise term with mean 〈*η*(*t*)〉 = 0, variance 〈*η*(*t*)^2^〉 = 1 and auto-correlation 〈*η*(*t*)*η*(*t*′)〉 that vanishes on timescales longer than 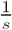. Thus, on timescales longer than 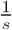, the allele frequency trajectory is just like that of a neutral mutation in a population of smaller size *Ne*^−*λ*^. On shorter timescales, the allele frequency trajectory will be more correlated in time than the frequency trajectory of a neutral population in a population of that size and will appear smoother. However, since large frequency changes of alleles at these frequencies will only occur on a timescale of order *Ne^−*λ*^ f*, which is much longer than 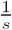, this description will be sufficient for describing site frequency spectra.

We emphasize that this coarse-grained model relies on the overall fluctuations in the fitness distribution of the population being negligible on relevant timescales, so that the average number of deleterious mutations per individual, 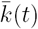 is approximately equal to *λ* (and crucially, independent of *f*). We expect that this approximation is valid when *N se*^−*λ*^ ≫ 1, because the overall fluctuations in the fitness distribution are small compared to *λ* Ne-her and Shraiman (2012). However, it is less clear whether this approximation continues to be appropriate as *N se*^−*λ*^ approaches more moderate, 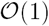 values. A more detailed exploration of these effects would require a path integral approach similar to that of Neher and Shraiman (2012), and is beyond the scope of this work.

### The trajectories of high frequency alleles, 1 – *f* ≪ 1

The coarse grained neutral model from the previous section breaks down when the allele frequency of the mutant exceeds 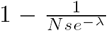. These total allele frequencies are attained when the frequency of the founding genotype *f*_0_ exceeds the frequency 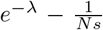. Mutant lineages that reach such high frequencies are almost certain to fix in the 0-class. Once this happens, all individuals that carry the wild type allele at the locus will also be linked to a deleterious variant. Thus, though the mutant carries no inherent fitness benefit, it will thereafter appear fitter than the wild type, because it has fixed among the most fit individuals in the population. The mutant will therefore proceed to perform a true selective sweep and will drive the wild type allele to extinction.

At these high allele frequencies, 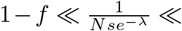 1, we can once again use the coupled branching process in Eq. 13 to describe the allele frequency trajectory of the wild type, *f*_wt_(*t*) = 1 – *f*(*t*). Seen from the point of view of the wild type, the fixation phase of the mutant corresponds to the extinction phase of the wild type. To obtain a description of the allele frequency trajectory of the wild type at these times, we can expand the generating function in Eq. 18 at long times, which yields

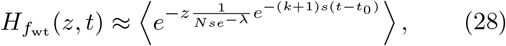

for some choice of *t*_0_ (see Appendix E for details). Note that, as before, Eq. 28 is valid only as long as 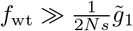 (i.e. as long as the size of the lineage in the 1-class exceeds 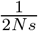). Once the frequency of the wild type in the 1-class falls below 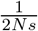, we can no longer treat this class deterministically. Once this happens, the part of the wild type that is in the 1-class will drift to extinction within about 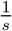 generations, whereas its bulk will continue to decay at a rate proportional to its average fitness, –2*s*. This will go on for as long as the frequency in the 2-class is larger than 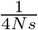, corresponding to the total frequency of the lineage being larger than 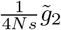. Once the frequency of the wild type in the 2-class also falls below 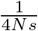, the bulk of the lineage will continue to decay even more rapidly, at rate –3s, and so on.

Thus, the wild type goes extinct in a staggered fashion, dying out in classes of higher fitness first, and declining in relative fitness in this process. As a result, the effective negative fitness of the wild type increases as its frequency declines, leading to an increasingly rapid exponential decay of the allele frequency (see right inset in Fig. 4A). The average fitness of the bulk of the wild type distribution is, to leading order, given by (see Appendix H)

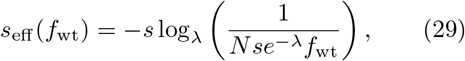

valid for 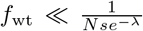, which means that the frequency trajectory of the wild type in this phase obeys

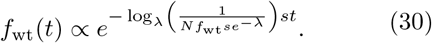

## THE SITE FREQUENCY SPECTRUM IN THE PRESENCE OF BACKGROUND SELECTION

Having obtained a distribution of allele frequency trajectories, we are now in a position to evaluate the site frequency spectrum. Since the trajectory of any lineage depends on the fitness of the background that it arose on, we will find it convenient to divide the total site frequency spectrum *p*(*f*) into the site frequency spectra of mutations with different ancestral background fitnesses, *p*(*f*, *k*). By definition, the total site frequency spectrum *p*(*f*) is the sum over these single-class frequency spectra,

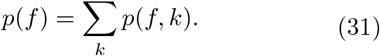

We evaluate the site frequency spectrum in three overlapping regimes, 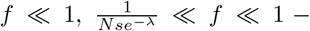 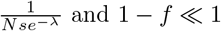

### The site frequency spectrum of rare alleles, *f* ≪ 1

The rare end of the frequency spectrum (f ≪ 1) consists of neutral alleles that (because they are rare) occurred on different genetic backgrounds. These alleles thus have independent allele frequency trajectories that can be described by the coupled branching process, Eq. 13. As long as 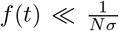, most lineage trajectories are dominated by genetic drift. Intuitively this result is simple: provided that the lineage is rare enough, selection pressures in any fitness class (or more precisely, the bulk of the fitness classes where the vast majority of such alleles arise) can be neglected compared to drift. Thus, the resulting site frequency spectrum is

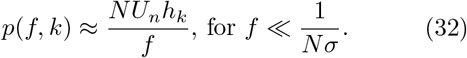

At these frequencies, the total site frequency spectrum is equal to

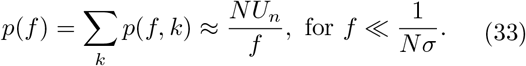

This agrees with our earlier intuition that at the lowest frequencies the entire population contributes to the site frequency spectrum, and also with the results of Wright-Fisher simulations (see Figure 5). Since the effects of selection are negligible, each fitness class contributes proportionally to its size, with the largest fitness classes contributing the most (see Figure 6). The deleterious mutation-free (*k* = 0) class has a negligible effect on the site frequency spectrum, contributing only a small proportion (proportional to its total frequency, *h*_0_ = *e*^−*λ*^) of all variants seen at these frequencies.

**FIG. 5.**
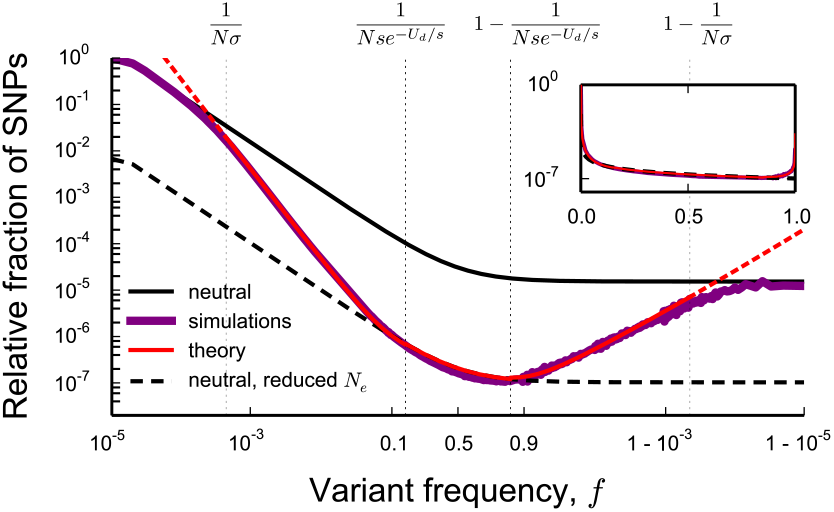
Comparison between the theoretical predictions for the site frequency spectrum (black, red) and Wright-Fisher simulations (purple). The simulated site frequency spectrum shown above is the same as the site frequency spectrum in Figure 1. At frequencies smaller than 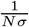 and larger than 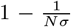, the theoretical predictions agree with the predictions for a neutral population with census size *N* (black line). Inside the range 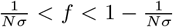, the theoretical predictions are given by the red line. The three different functions valid when 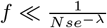 (Eq. 36), 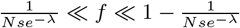 (Eq. 37) and 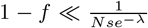 (Eq. 38) were joined using sigmoid functions 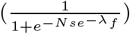. The logarithmic divergences at 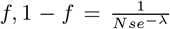 were controlled by adding small factors (0.5 and 1) to the logarithms in the denominators of Eq. 36 and Eq. 38, and then adjusting Eq. 36 and Eq. 38 using small multiplicative factors (equal to 2), see Appendix H.

**FIG. 6.**
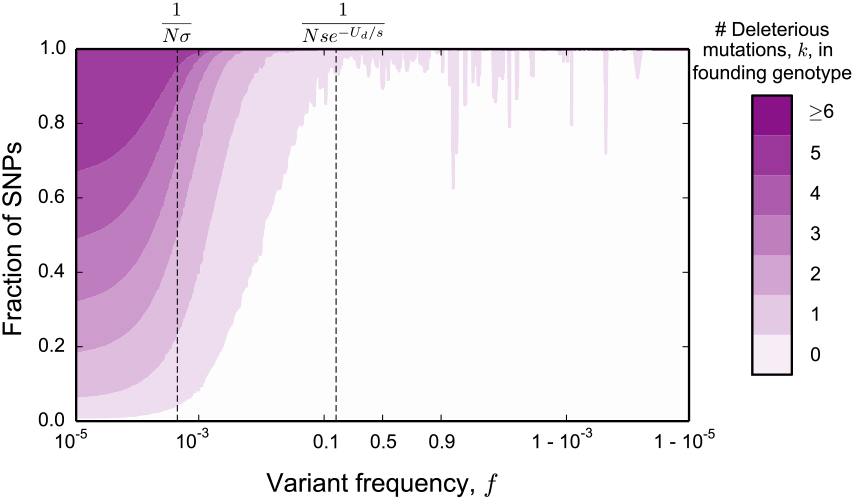
Proportions of polymorphisms at a given frequency that arose in genotypes that had a specified number of additional deleterious mutations compared to the most fit genotype at the locus at the time they arose. The parameters in these simulations are the same as the parameters used to generate site frequency spectra in Fig. 1 and Fig. 5. Note that for frequencies 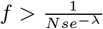, the entire site frequency spectrum is composed of lineages that arose due to neutral mutations in the *k* = 0 class (with only a few exceptions that arise due to rare ratchet events).

At larger frequencies, 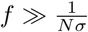, selection plays an important role in shaping allele trajectories and the site frequency spectrum. The overall contribution of mutations originating in class *k* near some frequency *f* is determined not only by the overall rate *NU*_*n*_*h*_*k*_ at which such mutations arise, but also by the probability that these mutations reach *f*, which declines with the initial deleterious load *k.* As a result, as *f* increases, the site frequency spectrum will become increasingly enriched for alleles arising in unusually fit backgrounds (see Figure 6).

The contributions to the site frequency spectra *p*(*f*, *k*) are straightforwardly obtained by integrating in time the distributions of allele frequency trajectories that we have described in the Analysis section,

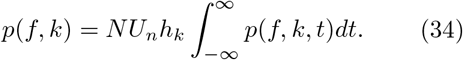

This integral is dominated by the peak phase of allele frequency trajectories, during which we have seen that the allele frequency is simply proportional to the weight in class 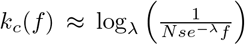, which corresponds to the class of lowest fitness in which the dynamics are not deterministic,

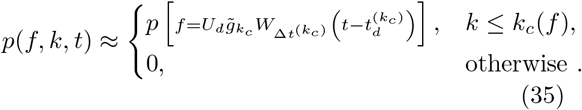

The overall site frequency spectrum is equal to the sum of these terms (Eq. 31). In Appendix H we show that this sum is well-approximated by the last term, corresponding to *k* = *k*_*c*_(*f*), and obtain that the site frequency spectrum in the rare end is, to leading order

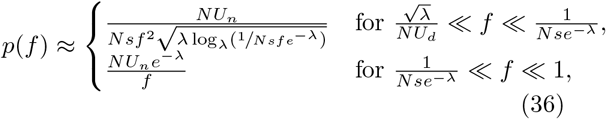

where the form of the frequency spectrum for *f* ≪ 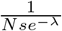 is valid up to a constant factor (see Appendix H for details). A comparison between these predictions and site frequency spectra obtained in Wright-Fisher simulations of the model is shown in Figure 5.

These results reproduce much of what we may have anticipated from our analysis of allele frequency trajectories. At frequencies 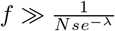, these peaks represent the mirrored and amplified trajectories in the mutation-free (*k* = 0) founding class. To reach these frequencies, mutations need to arise in the mutation-free class (which happens at rate *NU*_*n*_*e*^−*λ*^) and drift to substantial frequencies 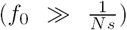. Since fluctuations in the founding class of lineages that exceed this frequency are slow compared to the timescale on which their deleterious descendants remain at their peak 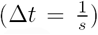, the entire allele frequency trajectory reproduces the fluctuations of the neutral founding class. Thus, neutral site frequency spectra proportional to *f*^−1^ emerge.

At smaller frequencies, 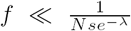, allele frequency trajectories reflect the smoothed out fluctuations in the high-fitness classes. At these frequencies, the site frequency spectrum is composed of a rapidly increasing number of polymorphisms for three reasons. First, lower frequencies correspond to smaller feeding class weights, which are more likely simply due to the effects of drift. Second, the number of individuals with *k* deleterious mutations in the locus increases with *k* (for *k* < *λ*), causing an increase in the overall rate at which alleles peaking at lower frequencies arise. This variation in the overall number of such alleles gives rise to the steeper power law, *f*^−2^, which can be compared to the distribution of peaks of individual lineages, which decays at most as *f*^−3/2^. Finally, peaks that occur at frequencies 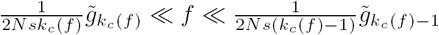 have duration of order 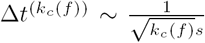, which declines with the frequency *f*, giving rise to the root logarithm factor.

### The site frequency spectrum at intermediate and high frequencies

At frequencies much larger than 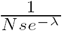 but still smaller than 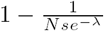, the allele frequency trajectory is described by an effective neutral model on coarse enough timescales 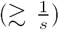. At these frequencies, the site frequency spectrum is

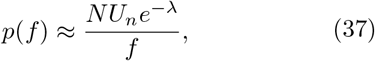

for

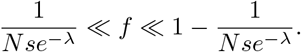

Note that this agrees with the result of the branching process calculation, which is valid in a part of this range, at frequencies corresponding to 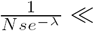 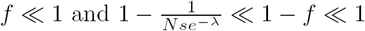.

This breaks down at even higher frequencies 1 – 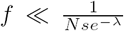. These frequencies correspond to the extinction phase of the wild type, during which the allele frequency no longer mirrors the frequency in the 0-class, but instead declines exponentially at an accelerating rate (see Eq. 30). Equation 30 can be straightforwardly integrated in time (see Appendix H for details), which yields the form of the site frequency spectrum at these high frequencies

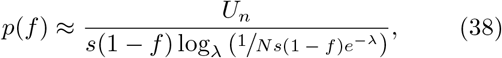

for

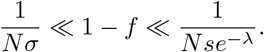

Finally, once the wild type frequency falls below 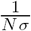, it will be in an analogous situation as the mutant at very low frequencies: independent of how it is distributed among the fitness classes, its trajectory will be dominated by drift, since most individuals in the population have fitness that does not differ from the mean fitness by more than *σ.* Thus, at these frequencies, the site frequency spectrum will once again agree with the site frequency spectrum of neutral loci unlinked to any selected sites in the genome

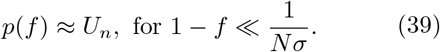

By comparing our predictions to the results of Wright-Fisher simulations, we can see that this argument correctly predicts the form of the site frequency spectrum in these regimes, as well as the frequencies at which these transitions happen (see Figure 5).

### The deleterious site frequency spectrum

So far, we have focused primarily on describing the trajectories and site frequency spectra of neutral mutations. However, because the trajectory of a neutral mutation that arises in an individual with *k* deleterious mutations is equivalent to the trajectory of a deleterious mutation that arises in an individual with *k* – 1 deleterious mutations, descriptions of trajectories of deleterious mutations follow without modification from our descriptions of trajectories of neutral mutations. The deleterious site frequency spectrum can thus be constructed from the single-class site frequency spectra of neutral mutations by a simple modification of the total rates at which new deleterious mutations arise (specifically, with the contribution to the deleterious site frequency spectrum of mutations arising in class *k* being equal to 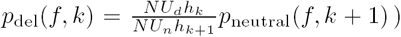. By summing these contributions, we find that the deleterious site frequency spectrum is to leading order

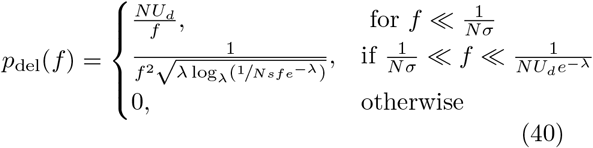

where the form proportional to 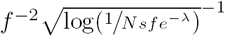 is once again valid up to a constant factor (see Appendix H).

### DISTRIBUTIONS OF EFFECT SIZES

The model we have thus far considered assumes that all deleterious mutations have the same effect on fitness, *s.* In reality, different deleterious mutations will have different fitness effects. In Appendix I, we show that as long as the variation in the distribution of fitness effects (DFE) is small enough that 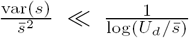, the effects of background selection are well-captured by a single-*s* model. In practice, this means that when considering moderate values of 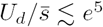, fractional differences in selection coefficients up to 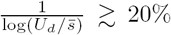 will not substantively alter allele frequency trajectories. In this case, the combined effects of these mutations are well-described by our single-s model.

However, when mutational effect sizes vary over multiple orders of magnitude, properties of the DFE will have an important impact on the quantitative details of the mutational trajectories that are not captured by our single-s model. The qualitative properties of allele frequency trajectories will remain the same (see Appendix I): alleles arising on unusually fit backgrounds will rapidly spread through the fitness distribution, peak for a finite amount of time about *t*_*d*_ generations later, and then proceed to go extinct at a rate proportional to their average fitness cost. However, the quantitative aspects of these trajectories will be different. For instance, small differences in the fitness effects of mutations 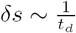 that do not impact the early stages of trajectories will be revealed on timescales of order *t*_*d*_, and affect the size and the width of the peak of the allele frequency trajectory. We have seen that these two quantities play an important role in determining the properties of allele frequency trajectories and the site frequency spectrum.

As a result of the fact that weaker effects and smaller differences in effect sizes play a more important role in later parts of the allele frequency trajectory, the DFE relevant during the early phases of the trajectory may be different than the DFE relevant in the later phases of the trajectory. Furthermore, since longer-lived trajectories are also those reach higher frequencies (having originated in backgrounds of higher fitness), this can result in a different DFE that is relevant at larger frequencies, compared to the DFE relevant at lower frequencies. As a result, it is possible that for certain DFEs, no single 'effective' effect size can be used to describe the trajectories at all frequencies. The full analysis of a model of background selection in which mutational effect sizes come from a broad distribution remains an interesting avenue for future work.

### DISCUSSION

In this work, we have analyzed how linked purifying selection changes patterns of neutral genetic diversity in a process known as background selection. We have found that whenever background selection reduces neutral genetic diversity, it also leads to significant distortions in the neutral site frequency spectrum that cannot be explained by a simple reduction in effective population size (see Figure 1). These distortions become increasingly important in larger samples and have more limited effects in smaller samples (Fig. 1B,C). In this sense, the sample size represents a crucial parameter in populations experiencing background selection.

By introducing a forward-time analysis of the trajectories of individual alleles in a fully linked genetic locus experiencing neutral and strongly selected deleterious mutations, we derived analytical formulas for the whole-population site frequency spectrum. These results can be used to calculate any diversity statistic based on the site frequency spectrum in samples of arbitrary size. Our results also offer intuitive explanations of the dynamics that underlie these distortions, and give simple analytical conditions that predict when such distortions occur. In addition to single-timepoint statistics such as the site-frequency spectrum, our analysis also yields time-dependent trajectories of alleles. We suggest that these may be crucial for distinguishing between evolutionary models that may remain indistinguishable based on site frequency spectra alone. We discuss these findings in turn below.

In the presence of strongly selected deleterious mutations 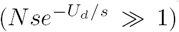, the magnitude of the effects of background selection critically depends on the ratio of the deleterious mutation rate, *U*_*d*_, to the selective cost of each deleterious mutation, *s.* Whenever *U*_*d*_ < *s*, both the overall genetic diversity and the full neutral site frequency spectrum *p*(*f*) are unaffected by background selection and *p*(*f*) is to leading order equal to

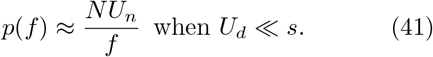

However, we have shown that the neutral site frequency spectrum follows a very different form when *U*_*d*_ ≫ *s:*

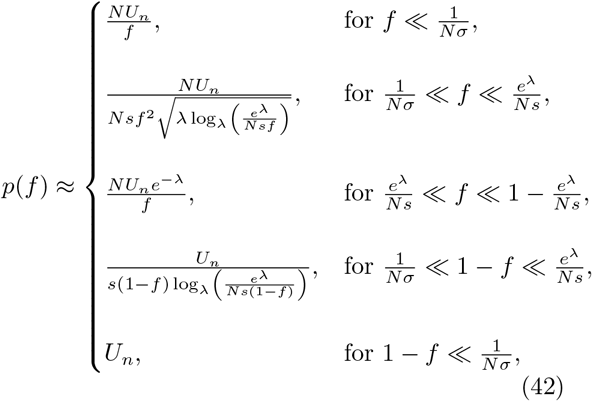

where 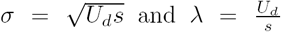. We note that *p*(*f*) matches the site frequency spectrum of a neutral population with a smaller effective population size *N*_*e*_ = *Ne*^−*λ*^ between derived allele frequencies 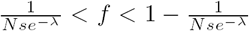, but deviates strongly outside this frequency range. This implies that summary statistics based on the site frequency spectrum (e.g. the average minor allele frequency) will start to deviate from the neutral expectation in samples larger than *Nse*^−*λ*^ = *“N_e_s”* individuals, but not in smaller samples (Fig. 1B,C).

Our approach is similar in spirit to the analysis of the distribution of frequencies in a general diffusion process by Ewens (1963), and to the analysis by Sawyer and Hartl (1992) of the frequency spectrum at an unlinked selected locus. However, we explicitly incorporate linkage between multiple selected sites. Such linkage is necessary for background selection to affect frequency spectra. However, linkage between multiple selected sites also complicates the trajectories of individual polymorphisms, which now depend not only on their own fitness effects but also on the effects of all other mutations that occur in the population.

Our analysis offers intuition into why background selection has an impact on site frequency spectra when *U*_*d*_ ≫ *s*, but not when *U*_*d*_ < *s.* When *U*_*d*_ ≫ *s*, a large majority of individuals in the population will carry some deleterious mutations at the locus, which results in substantial fitness variation within the population. This variation in fitness leads to a distinction between the absolute and relative fitness of an allele: although genotypes that carry *any* strongly deleterious variants are guaranteed to be eventually purged from the population, those that contain *fewer than average* deleterious mutations are still positively selected on shorter timescales. This results in strong non-neutral features in the frequency trajectories of alleles that arise on backgrounds that contain fewer than average deleterious mutations. Their trajectories are characterized by rapid initial expansions, followed by a peak, and eventual exponential decline (Figure 2). These deterministic aspects of allele frequency trajectories are similar to those seen by Neher and Shraiman (2011) in models of linked selection in large facultatively sexual populations.

The fluctuations in these allele frequency trajectories are primarily driven by the fluctuations in the numbers of most-fit individuals that carry the allele. This is closely related to the fluctuations in the population fitness distribution studied by Neher and Shraiman (2012) in an analysis of Muller's ratchet. Here we have quantified how these fluctuations propagate to shape the statistics of allele frequency trajectories, finding that fluctuations in the number of most-fit individuals that happen on a timescale shorter than 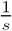 are smoothed out due to the finite timescale on which selection can respond. In contrast, fluctuations that happen on timescales longer than 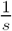 are faithfully reproduced in the allele frequency trajectory. This smoothing of fluctuations on a finite timescale introduces an additional fundamentally non-neutral feature in the total allele frequency trajectory, which distorts the site frequency spectrum at frequencies below 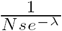.

### The frequency of a mutation tells us about its history and future

In addition to describing the expected site frequency spectrum at a single time point, our analysis of allele frequency trajectories allows us to calculate time-dependent quantities such as the posterior distribution of the past frequency trajectory of polymorphisms seen at a particular frequency, their ages, and their future behavior. For example, since the maximal frequency a mutation can attain strongly depends on the fitness of the background in which it arose (with lower-fitness backgrounds constraining trajectories to lower frequencies), observing an allele at a given frequency places a lower bound on the fitness of the background on which it arose. This in turn is informative about its past frequency trajectory. For example, alleles observed at frequencies 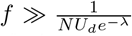 almost certainly arose in an individual that was among the most fit individuals in the population and experienced a rapid initial exponential expansion at rate *U*_*d*_, while alleles observed at frequencies 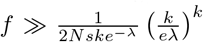 very likely arose on backgrounds with fewer than *k* deleterious mutations compared to the most fit individual at the time. We emphasize that these thresholds are substantially smaller than the naïve thresholds obtained by assuming that a mutation arising on a background with *k* mutations can only reach the ‘drift barrier’ 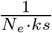 corresponding to unlinked deleterious of locus of fitness *ks* in a population of effective size *N*_*e*_ = *Ne*^−*λ*^.

The fitness of the ancestral background that a mutation arose on is not only interesting in terms of characterizing the history of a mutation, but is also informative of its future behavior. In the strong selection limit of background selection that we have considered here (*Nse*^−*λ*^ ≫ 1), deleterious mutations can fix in the population only exponentially rarely (Neher and Shraiman, 2012). Thus, mutations arising on backgrounds already carrying deleterious mutations must eventually go extinct. We have shown that the site frequency spectrum at frequencies 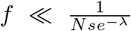 is dominated by mutations arising on deleterious backgrounds. Furthermore, we have shown that most polymorphisms seen at these frequencies are at the peak of their frequency trajectory. This means that we expect the frequency of such polymorphisms to decline on average. Thus, if we were to observe the population at some later timepoint, we expect that the polymorphisms present at such low frequencies should on average be observed at a lower frequency. In Figure 7 we show how the average change in frequency after *Ne*^−*λ*^ generations depends on the original frequency that a mutation was sampled at, *f*. Note that the expectation for a neutral population of any size is that the average allele frequency change is exactly equal to zero. In the presence of background selection, this is no longer true for neutral mutations previously observed at frequencies 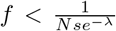 and 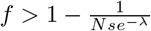 (see Fig. 7).

**FIG. 7.**
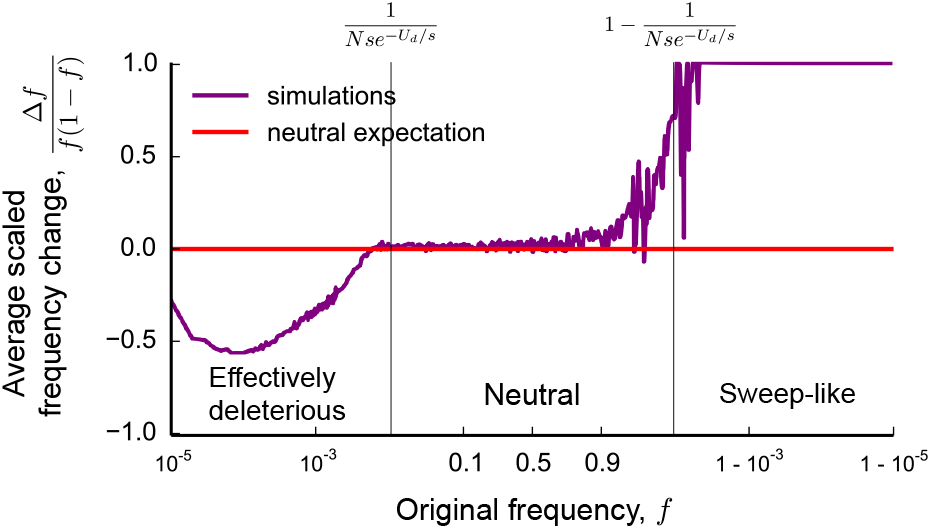
The average change in frequency of an allele observed at frequency *f* in an earlier sample. The pairs of frequencies were sampled in two consecutive sampling steps in Wright-Fisher simulations in which *N* = *10*^5^, *NU*_*d*_ = 4*Ns* = 2 · 10^4^. In the first step, the frequencies of all polymorphisms in the population and their unique identifiers were recorded. In the second sample, which was taken 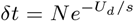 generations later, the frequencies of all polymorphisms seen in the first sample were recorded (if a polymorphism had gone extinct, or fixed, a frequency *f*(*δt*) = 0, or 1, was recorded). The average represents the average frequency change over many distinct polymorphisms, and the curve has been smoothed using a Gaussian kernel with width < 5% of the minor allele frequency.

In contrast, since polymorphisms observed in the range 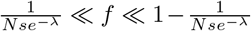 must have originated in a mutation-free background, and since their dynamics reflects neutral evolution in this 0-class, the overall dynamics of such alleles are neutral. Therefore, though drift will lead to variation in the outcomes of individual alleles in this range, the average expected frequency change is equal to zero. This expectation is confirmed by simulations (Fig. 7).

Finally, we have seen that polymorphisms seen at frequencies 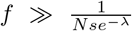 will typically already have replaced the wild type allele within the 0-class. Thus, the wild-type allele must eventually go extinct (except for exponentially rare ratchet events). In other words, polymorphisms seen at these frequencies are certain to fix, replacing the ancestral allele at some later point in time (see Fig. 7). Together, these results show that the site frequency spectrum can be divided into three regimes, in which the dynamics of individual neutral alleles are effectively negatively selected, effectively neutral, and effectively positively selected. These effective selection pressures arise indirectly, as a result of the fitnesses of the variants a neutral mutation at a given frequency is likely to be linked to.

### The distinguishability of models based on site frequency spectra

As has long been appreciated, background selection can lead signatures in the site frequency spectrum that are qualitatively similar to population expansions and selective sweeps (Charlesworth et al., 1993, 1995; Good et al., 2014; Gordo et al., 2002; Hudson and Kaplan, 1994; Nicolaisen and Desai, 2012; O’Fallon et al., 2010; Tachida, 2000; Walczak et al., 2011; Williamson and Orive, 2002). Here we have shown that these similarities are not only qualitative, but (up to logarithmic corrections) also quantitatively agree with the site frequency spectra produced under these very different scenarios (Lea and Coulson, 1949; Mandelbrot, 1974; Yule, 1924). This suggests that distinguishing between these models based on site frequency spectra alone may not be possible. We emphasize that these effects of background selection that mimic population expansions are seen in *neutral* site frequency spectra in a model in which the population size is fixed, so using synonymous site frequency spectra to ‘correct’ for the effects of demography may not always be justified.

The quantitative agreement between the effects of background selection and positive selection that we have seen in the high frequency end of the frequency spectrum is not purely incidental. In the presence of substantial variation in fitness, alleles that fix among the most fit genotypes in the population are in a sense truly positively selected, because they are linked to fewer deleterious mutations than average. As a result, sweep-like behaviors can occur in the absence of positive selection, as long as there is substantial fitness variation, independent of the source of this variation (i.e. whether it arose as a result of beneficial or deleterious mutations). In this case, these models may be indistinguishable even using time-resolved statistics, because the allele frequency trajectories themselves have similar features.

In other cases, time-resolved statistics may be able to differentiate between models that produce similar site frequency spectra. For example, under background selection the low frequency end of the site frequency spectrum is dominated by mutations that are linked to a larger than average number of deleterious variants; alleles in this regime are therefore expected to decline in frequency on sufficiently long timescales (of order 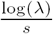. In contrast, in an exponentially expanding population mutations present at these frequencies are very unlikely to change in frequency during the expansion. Thus we may be able to distinguish between these models using samples from the same population spaced far enough apart in time.

### Relationship to results on weakly selected deleterious mutations

Throughout this work, we have assumed that selection against deleterious mutations is strong (i.e. *Nse*^−*λ*^ ≫ 1), such that they are exponentially unlikely to fix (i.e. Muller's ratchet is rare). In the opposite case where selection is weak enough that deleterious mutations have a substantial probability of fixation (which occurs when *Nse^−*λ*^ ≲* 1), the population rapidly ratchets to lower fitness at the locus. In this case, earlier work has shown that genealogies approach the Bolthausen-Sznitman coalescent when the fitness variance, *σ*, in the population is sufficiently large, *Nσ ≫* 1 (Good et al., 2014; Kosheleva and Desai, 2013; Neher and Hallatschek, 2013; Neher and Shraiman, 2011). In this case, the resulting site frequency spectrum scales as *f*^−2^ at low frequencies, and as [(1 − *f*)log(^1^/(1 −*f*))]^−1^ at high frequencies (Neher and Hallatschek, 2013). These forms of the site frequency spectrum differ from our results for populations with large fitness variance by the large root logarithm factor at low frequencies and by a large constant factor proportional to log(*Nse*^−*λ*^) at high frequencies, as well as by the absence of the effectively neutral regime at intermediate frequencies. As *N se*^−*λ*^ approaches 1, this intermediate regime disappears and the site frequency spectrum starts approaching the Bolthausen-Sznitman limit (though see Good et al. (2014) for a discussion of weakly selected mutations and moderate *Nσ*).

Recently, Hallatschek (2017) has studied allele frequency trajectories that arise in the forward-time dual of the Bolthausen-Sznitman coalescent. Our analysis reveals many of the interesting features seen in that work. For instance, we have seen that once an allele spreads through the fitness distribution and reaches mutation-selection balance, an effective frequency-dependent selection coefficient emerges *s*_eff_(*f*),

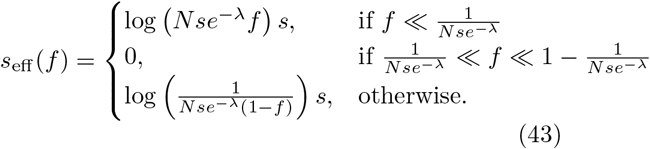

This effective selection coefficient arises due to the deleterious mutations that neutral mutations are linked to, and changes with the frequency *f* of the mutation as high-fitness individuals within the neutral lineage drift to extinction. This is analogous to the fictitious selection coefficient, *s*_fic_(*f*) = 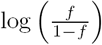, that emerges in the model analyzed by Hallatschek (2017). The difference in the form of the effective selection coefficient between our results and the Hallatschek (2017) model is large when *Nse*^−*λ*^ ≫ 1, but becomes negligible as *Nse*^−*λ*^ → 1; it underlies the differences between the site frequency spectra of rapidly adapting or ratcheting populations and the strong background selection limit that we have considered here.

### Extensions and limitations of our analysis

We have studied a simple model of a perfectly linked locus at which all mutations are either neutral or deleterious with the same effect on fitness, *s.* Our primary goal has been to describe the qualitative and quantitative effects of background selection on frequency trajectories and the site frequency spectrum within this simplest possible context. However, it is important to note that the assumptions of our model are likely to be violated in natural populations. In many cases, these additional complications do not change the general conclusions of our analysis. For example, the qualitative properties of the trajectories and site-frequency spectra described here apply when deleterious mutations have a broader distribution of effect sizes, and we have shown here that our results are quantitatively unchanged when the distribution (DFE) of effect sizes is sufficiently narrow 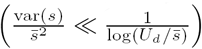. On the other hand, when the DFE is very broad, additional work will be required to determine the quantitative properties of site frequency spectra. We anticipate different parts of the DFE may be important at different frequencies in sufficiently broad DFEs. If this is true, this would be an unusual feature of strong negative selection that does not arise in the case of strong positive selection, in which the effects of DFEs can usually be summarized by a single, predominant fitness effect (Good et al., 2012).

Finally, our assumption of perfect linkage in the genomic segment is likely to be violated in sexual populations, in which sites that are separated by shorter genomic distances are more tightly linked than distant sites. However, even in the presence of recombination, alleles will remain effectively asexual on short enough genomic distances, and are effectively freely recombining on long enough genomic distances (Franklin and Lewontin, 1970; Slatkin, 1972). In this case, a standard heuristic is to treat the genome as composed of freely recombining asexual blocks. This heuristic has been shown to yield a rough approximation to diversity statistics when the ‘effective block length’ is set by the condition that each block typically recombines once on the timescale of coalescence, both in the case of background selection, as well as in the case of rapid adaptation (Good et al., 2014; Neher et al., 2013; Weissman and Hallatschek, 2014).

However, our analysis highlights that many of the interesting features of allele frequency trajectories in the presence of background selection occur on timescales much shorter than the timescale of coalescence. On these timescales, alleles will be fully linked on much longer genomic distances than this effective block length. This effect will be particularly important for young alleles, which are linked to long haplotypes because of the limited amount of time that recombination has had to break them up. On longer timescales, the length of the genomic segments that these alleles are linked to will become progessively shorter, but will typically not fall below the ‘effective block size’ on any timescale. Given the strong dependence of allele frequency trajectories on the total mutation rate along this segment, it is less clear what effect such linkage to increasingly shorter genomic segments has on the statistics of allele frequency trajectories. A more detailed analysis of the effects of background selection in linear genomes remains an intriguing direction for future work.

## ACKNOWLEDGEMENTS

We thank Oskar Hallatschek, Joachim Hermisson, Katherine Lawrence, Matthew Melissa, Richard Neher, Daniel Rice, Boris Shraiman, Shamil Sunyaev and John Wakeley and the members of the Desai lab for useful discussions and helpful comments on the manuscript. Simulations in this article were run on the Odyssey cluster supported by the FAS Division of Science Research Computing Group at Harvard University. This work was supported in part by the Simons Foundation (grant 376196), grant DEB-1655960 from the NSF, and grant GM104239 from the NIH. The authors also acknowledge the Kavli Institute for Theoretical Physics at UCSB, supported in part by the NSF grant PHY-1125915, NIH grant R25GM067110, and the Gordon and Betty Moore Foundation grant 2919.01.

## Appendix A: The propagation of fluctuations in the size of the founding class

In this Appendix, we consider in more detail how fluctuations in the size of the lineage in the founding class propagate to affect the total allele frequency. For this purpose, it will be convenient to consider a neutral mutation that has arisen in the *k* = 0 class sufficiently long ago that it is in mutation-selection balance. Let the total frequency of the lineage be *f*. As in the main text, we denote the frequency of the part of the lineage that is in class *i* by *f*_*i*_. In mutation-selection balance, the *f*_*i*_ will satisfy

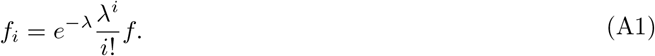

Consider what happens if the frequency *f*_0_ of the founding genotype changes suddenly to some value *f*_0_ + *δf*_0_. Based on the deterministic solution, after a time *t*, this will lead to a change in the frequency of the part of the lineage in class *i*, *δf*_*i*_(*t*), of

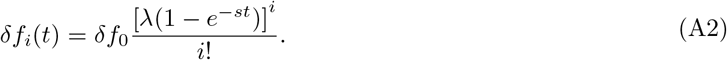

In other words, the relative change in the frequency of the lineage in the *i*-class is

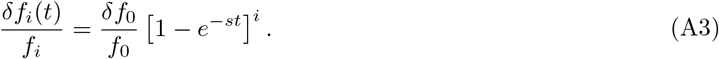

This approaches 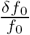 at long times as the allele re-establishes mutation-selection balance. However, we can see from Eq. (A3) that this change is not felt at the same time in all classes. In the 1-class, the frequency changes gradually, at rate *s* (Eq. (A3)), and results in a proportional change roughly 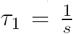 generations later. In general, in the *i*-class, this change is felt after a total delay of roughly 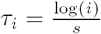 generations. Thus, the change propagates from class *i* to class *i* + 1 over the course of

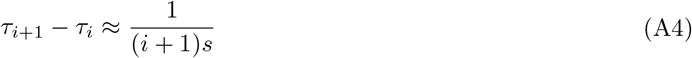

generations.

Ultimately, 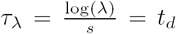 generations later, this change will have been felt in a substantial fraction of the fitness distribution. Fitness classes near the mean of the distribution (which is *λ* classes below the 0-class) are those that exhibit the largest absolute change in frequency, since they contain the largest number of individuals when the lineage is in mutation-selection balance. Thus, changes in these classes account for a large proportion of the change in the total allele frequency, which explains the origin of the delay timescale, *t*_*d*_, that we have introduced in the main text.

## Appendix B: The large deviations from average behavior caused by genetic drift

In this Appendix, we consider the importance of drift in each individual fitness class on the overall allele frequency. In the first subsection, we revisit a standard argument to explain why fluctuations due to genetic drift in the frequency of the founding genotype can never be neglected, framing it in terms that will be useful when considering the importance of drift in classes below the founding class. In the next subsection, we build on this argument to explain why the effects of drift become negligible in all classes *i* in which the frequency of the component of the lineage in that class, 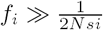, but cannot be neglected in all classes in which the frequency does not exceed that threshold.

### 1. The importance of genetic drift in the founding class

The essential reason why drift can never be neglected in the early phase of a trajectory is that deviations from the low frequency average behavior caused by drift are not small perturbations, but are extremely broadly distributed. Consider for instance a mutation that arises in class *k.* As we explain in the main text, the founding genotype feels an effective selection coefficient equal to −*ks.* The deterministic trajectory of the founding genotype is therefore

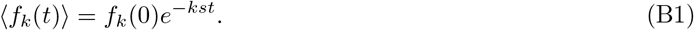

In other words, the ‘deterministic trajectory’ of a neutral founding genotype (*k* = 0) is a flat line, whereas the deterministic trajectory of a deleterious founding genotype (*k* > 0) decays exponentially at rate *ks.*

However, we know that drift leads to large deviations from the deterministic behavior in Eq. (B1). In fact, we have mentioned that when 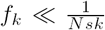, drift can lead to an *x*-fold increase above this expectation with probability 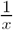 (Fisher, 2007). Thus, the deviations from the deterministic expectation due to drift are distributed according to an extremely broad power law. As a result, large deviations from Eq. (B1) are very likely. For lineages arising in the 0-class, these deviations can take the frequency of a lineage all the way to fixation. However, deleterious founding genotypes with *k* > 0 are exponentially unlikely to exceed the drift barrier at 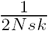. Thus, the distribution of deviations from the mean, deterministic behavior of these founding genotypes also follows the same power law at low frequencies 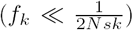, but is capped by selection at frequencies exceeding 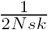. As a result, the effects of drift on trajectories of deleterious mutations become perturbative at sufficiently large frequencies and can therefore be neglected when 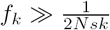.

Because fluctuations in *f*_*k*_ always propagate to classes of lower fitness, drift in the founding class has an important impact on the overall allele frequency whenever it has an important impact on *f*_*k*_. This means that the overall frequency trajectory of alleles founded in the 0-class will always be affected by drift in *f*_*k*_, which will cause large power law distributed deviations from the deterministic expectation of the *total* allele frequency trajectory. Similarly, the overall frequency trajectory of alleles founded in a class with *k* > 0 deleterious mutations will be dominated by drift at low frequencies 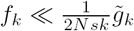, but capped from exceeding frequencies larger than 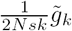 (see also Appendix E).

### 2. The importance of genetic drift in classes below the founding class

Given these arguments, one may wonder whether the effects of drift are also important in classes below the founding class, with *i* > *k* deleterious mutations. Deviations from deterministic behavior in these classes (i.e. in *f*_*i*_) are also propagated to classes of lower fitness. Such deviations in *f*_*i*_(*t*), if large, will also have a large impact on the overall frequency trajectory of the allele, *f*(*t*). However, since classes below the founding class receive substantial mutational input from higher classes, it is not immediately clear whether the effects of drift on *f*_*i*_(*t*) will 'average out' as a result of these mutations, or whether drift can still lead to large deviations from the deterministic expectation for *f*_*i*_(*t*). In Appendix E we show by formally analyzing the distribution of allele frequency trajectories that drift in class *i* is negligible when 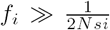, and in this Appendix we give a heuristic argument explaining why this threshold arises.

The threshold 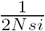 is reminiscent of the drift barrier relevant for single deleterious loci of fitness *is.* However its relevance in classes below the founding class is not immediately obvious. Although the individuals in class *i* also feel an effective selection pressure equal to *–is*, new mutational events from class *i* – 1 counter these effects of selection. Thus, it is not obvious that the combination of the opposing effects of mutation into the class and selection within the class will be stronger than the effects of drift whenever 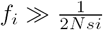 (as opposed to some other threshold that also depends on *U*_*d*_*f*_*i−1*_).

To gain insight into this, we consider in more detail the effects of individual mutational events into class *i.* Each of these mutational events can be thought of as founding a new ‘sub-lineage’ in class *i.* The frequency trajectory of each sub-lineage is the same as that of a single locus with fitness *−is* and the overall trajectory *f*_*i*_(*t*) is equal to the sum of the trajectories of these sub-lineages. When a sub-lineage is small, drift will lead to large deviations from its average (deterministic) frequency trajectory, which is also given by Eq. (B1). However, as in the founding class, at frequencies larger than 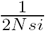, these deviations are capped by the effects of selection. Thus, the drift barrier 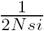 represents the frequency above which fluctuations cannot lead to large deviations of individual sub-lineages from the average behavior.

To understand when drift has an important impact on the overall *i*-class trajectory, *f*_*i*_(*t*), we can consider how these deviations in the trajectories of the sub-lineages add. At sufficiently small frequencies 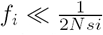, the overall trajectory *f*_*i*_ will be equal to the sum of random trajectories that have an extremely broad distribution. In this case, the sum will be dominated by the trajectory of the largest sub-lineage, which will be very different than the average trajectory. Thus, even when the total number of mutational events into class *i* is large, the effects of genetic drift in class *i* may not be negligible if each of these mutational events results in a relatively small trajectory. In other words, fluctuations due to drift in the frequency trajectories do not 'average out', but are rather dominated by the largest deviation from the mean. Conversely, when the total number of sub-lineages is large enough that many of them reach the frequency 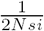 (which is guaranteed to happen if the total number of mutational events into the *i*-class is much larger than 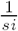), the overall frequency of the lineage will be much larger than 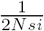. In this case; the largest event is no longer be very different than the average event; the effects of genetic drift are therefore negligible compared to the effects of selection. The transition between these two behaviors happens when 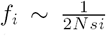, which roughly corresponds to exactly one sub-lineage exceeding 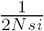. We discuss these effects using a more formal approach in Appendix G. Note that by extending this argument to classes *i* + 1 and lower, we can verify that once the frequency trajectory in class *i* exceeds 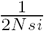 and becomes predominantly shaped by mutation and selection, the frequency of the allele in all lower fitness classes is also guaranteed to exceed the corresponding frequency thresholds, which is why we can also neglect the effects of drift in all classes below a class in which the frequency exceeds 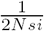.

## Appendix C: The generating function for the total size of the labelled lineage

In this Appendix, we consider the generating function for the total frequency of the lineage,

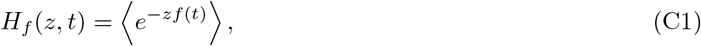

and derive a partial differential equation describing how it changes in time. As described in the main text, when the size of the lineage is small (*f*(*t*) ≪ 1), its dynamics are described by the coupled system of Langevin equations for the components *f*_*i*_(*t*) of the total frequency *f* that denote the frequency of the part of the lineage that carry *i* deleterious mutations,

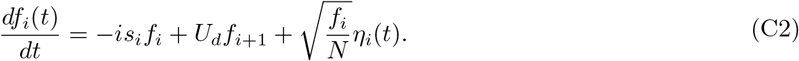

In Eq. (C2), the *η*_*i*_ are independent uncorrelated Gaussian noise terms. The total allele frequency is equal to the sum of these components, 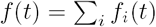.

Note that the total allele frequency *f*(*t*) is not a Markov random variable, since its evolution depends on the details of the distribution of the individuals within the lineage among the fitness classes. However, the frequencies of the components *f*_*i*_(*t*) are jointly Markov, with their joint distribution described by the joint generating function

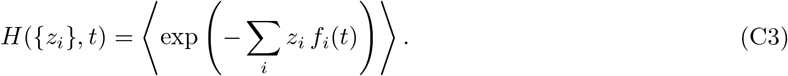

The generating function for *f*(*t*) can be obtained from the joint generating function by setting *z*_*i*_ = *z* for all *i.* We can obtain a partial differential equation for the joint generating function by Taylor expanding *H*({*zi*}, *t* + *dt*) and substituting in the differentials 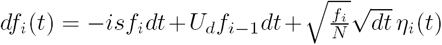 from Eq. (C2), which yields

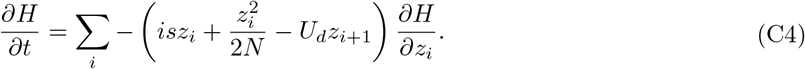

We can solve this PDE for the joint generating function by using the method of characteristics. The characteristic curves *Z*_*i*_(*t* – *t′*) are defined by

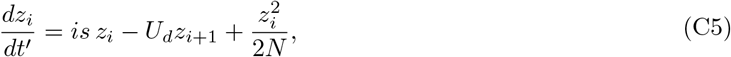

and satisfy the boundary condition *z*_*i*_(*t*) = *z.* The linear terms in the characteristic equation arise from selection and mutation into and out of the *i* class, and the nonlinear term arises from drift. Along these curves, the generating function is constant, and so 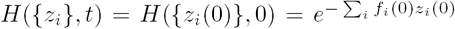, where the initial condition 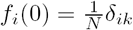 corresponds to a single individual present in class *k* at *t* = 0. Thus, to obtain a solution for the joint generating function we need to integrate along the characteristics in Eq. (C5) backwards in time from *t′* = 0 to *t′* = *t*. In the next few Appendices, we obtain these solutions in the limits of weak (*U*_*d*_ ≪ *s*) and strong mutation (*U*_*d*_ ≫ *s*).

## Appendix D: Trajectories in the presence of weak mutation (*U*_*d*_ ≪ *s*)

When deleterious mutations arise more slowly than selection removes them 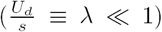, deleterious descendants of a lineage are much less numerous than the founding genotype. To see this, we can expand the characteristics *z*_*i*_(*t – t′*) in powers of the small parameter *λ*. At leading order, the characteristics are uncoupled and can be straightforwardly integrated to obtain

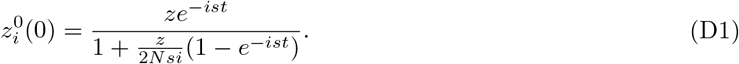

By substituting this zeroth order solution into Eq. (C5), we find that corrections due to deleterious descendants are 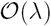, and are therefore small uniformly in *z.* Thus, the generating function for the total *f* of the labelled lineage *t* generations after arising in class *k* is

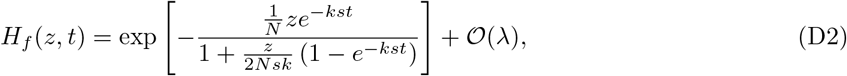

which agrees with classic results by Kendall (1948) for the generating function of independently segregating loci of fitness *−ks.*

Eq. (D2) can be inverted to obtain the probability distribution, *p*(*f*, *t*), by an inverse Laplace transform,

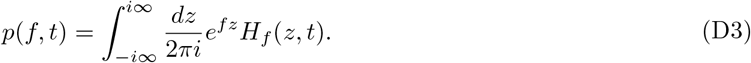

This distribution is well known, and can be obtained by standard methods. Noting that 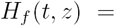 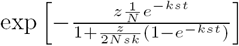 has a single essential singularity at 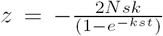, we can perform the integral above either exactly by contour integration (by closing the contour using a large semicircle in the left half-plane and a straightforward application of the residue theorem, which gives a solution in terms of Bessel functions), or approximately by the method of steepest descents (taking care to deform the contour to pass through the saddle point on the right of the essential singularity). By carrying out this inverse Laplace transform, we obtain that the extinction probability by time *t* is

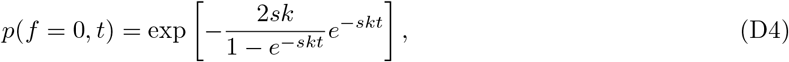

which becomes of order one when 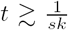, in agreement with our intuition that a lineage of fitness *ks* can only survive for order 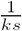 generations. For non-extinct lineages, the probability distribution of the frequency is

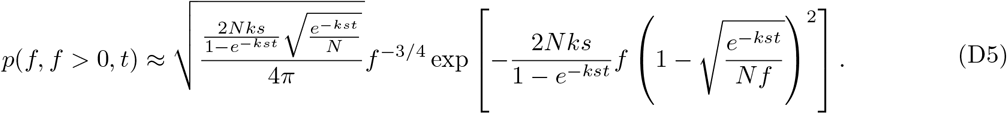

The site frequency spectrum can be obtained from this distribution of frequencies by integrating Eq. (D5) in time, or by an alternative method that we present in Appendix F.

## Appendix E: Trajectories in the presence of strong mutation (*Ud* ≫ *s*)

When deleterious mutations arise faster than selection can remove them, mutation will play an important role in shaping the trajectory. The relative strength of mutation and selection compared to drift will depend on the frequency of the lineage. Drift will remain the dominant force at frequencies 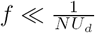. However, at larger frequencies, the mutation and selection terms will become important, and we will see that the effects of drift in classes of low enough fitness become negligible.

### 1. Small lineages 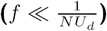

The dominant term in the characteristic equation in this regime (which corresponds to z ≫ *NU*_*d*_ in the generating function) is the drift term

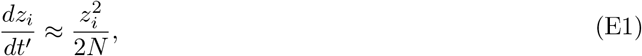

which has the solution

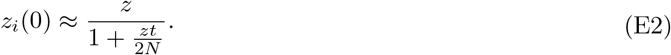

We can verify that mutation and selection are negligible compared to drift on timescales of order 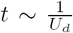 as long as *i* ≲ 2*λ*. Note that this condition (*i ≲ λ*) is satisfied for essentially all of the individuals in the population since 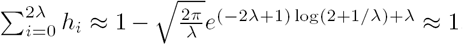. By summing the *Z*_*i*_(0) terms, we find that on these timescales the generating function for the frequency of the mutation is

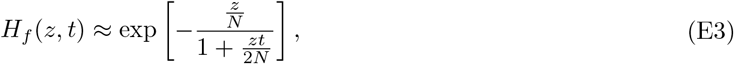

which is just the generating function for the frequency of a neutral lineage (cf. Eq. (D2)). On longer timescales 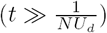, this approximation breaks down, and mutation and selection cannot be neglected for lineages arising in fitness classes far above the mean of the fitness distribution (with 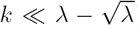). This is because the probability that a portion of the lineage in a class with fewer than 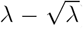 mutations has drifted to a high enough frequency to feel the effects of mutation and selection becomes substantial on longer timescales, which can also be seen from the probability distribution of non-extinct lineages (Eq. (D5)). We consider the generating function of these unusually fit mutations at these higher frequencies in the next subsection. In contrast, mutations that arise on more typical backgrounds with 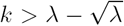 mutations can drift to higher frequencies, of order 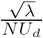, before feeling the effects of selection, but cannot substantially exceed a total frequency 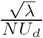. We analyze their trajectories in the following subsection.

### 2. Large lineages (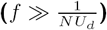) arising on unusually fit backgrounds 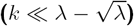

In lineages that reach higher frequencies, a large number of deleterious descendants arise every generation. This leads to strong couplings between the sizes of the components of the lineage in different fitness classes, and diminishes the importance of genetic drift in classes of lower fitness, which receive large numbers of deleterious descendants from classes of higher fitness. We find that in classes of low enough fitness, the effects of genetic drift are negligible and the dominant balance is between the linear mutation and selection terms.

The solution to the linear (deterministic) problem has been obtained by Etheridge et al. (2007), but we reproduce the derivation briefly for completeness. In the absence of drift, the characteristics evolve according to

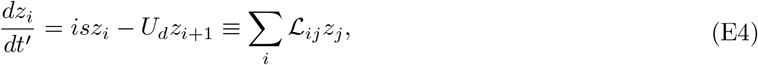

which defines the linear operator 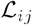. 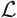 has right eigenvectors 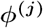 with eigenvalues – *js* given by and corresponding left eigenvectors

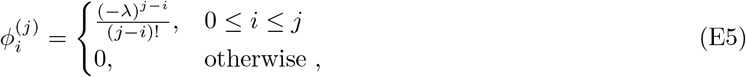

and corresponding left eigenvectors

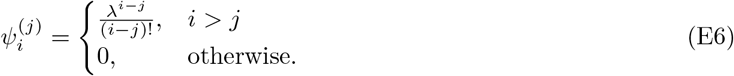

We can verify that the left and right eigenvectors are orthonormal 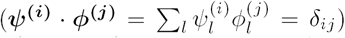. By eigenvalue decomposing *Z*_*i*_(*t*) and integrating backwards in time from *t′* = 0 to *t′* = *t*, we obtain *Z*_*i*_(*t* – *t*') = 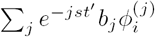, where the amplitudes *b*_*j*_ are set by the boundary condition at 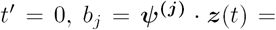 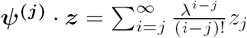. Finally, a summation yields

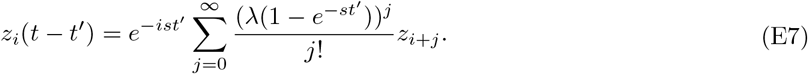

Setting the boundary condition at 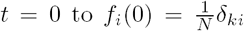 and evaluating *z*_*k*_(0), we reproduce the result by Etheridge et al. (2007): in the absence of genetic drift, the descendants of the labelled lineage follow a Poisson distribution that starts in class *k* and has mean *λ*(l – *e*^*–st*^) and amplitude 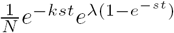.

To evaluate the effect of genetic drift on the total size of the lineage at some later time point we set *z*_*i*_ = *z*. A sufficient (but not necessary) condition for genetic drift in class *i* being negligible in determining the total size of the lineage at some later time point *t* is that the nonlinear term 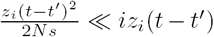 uniformly in *t′.* In the vicinity of some frequency *f*(*t*) ~ *f*, corresponding to 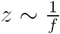, this will be true as long as

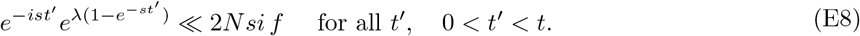

Since the left hand side is bounded by 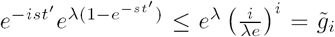, the inequality is guaranteed to be satisfied uniformly in *t* as long as

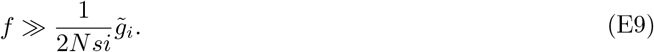

Defining *k*_*c*_(*f*) to be the largest integer for which 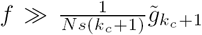, we can verify that genetic drift is negligible in all classes with *i* > *k*_*c*_(*f*) but not in class *k*_*c*_(*f*).

Note that self-consistency of the deterministic solution for *k*_*c*_ < *λ* implies that when 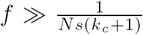, the frequency of the part of the allele in class *i* satisfies 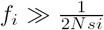 for all i > *k*_*c*_, but not for *i* ≤ *k*_*c*_. Also note that this inequality can only be satisfied for some *k*_*c*_ < *λ* if the founding class is sufficiently far above the fitness distribution 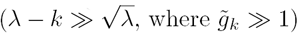. We return to lineages founded in classes with 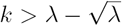 mutations in the next subsection.

Thus, since genetic drift has a negligible effect in classes containing more than *k*_*c*_ deleterious mutations, the characteristics *z*_*i*_(*t′*) are given by the deterministic solution above, which we have already integrated. The frequency of the part of the lineage in classes with *i* > *k*_*c*_ is therefore a deterministic function of the frequency trajectory in class *k*_*c*_, *f*_*k*_*c*__(*t*). We can solve for this deterministic function straightforwardly by explicitly including *f*_*k*_*c*__(*t*) as a variable mutational source term for classes of lower fitness. This yields an expression for the generating function of the entire lineage

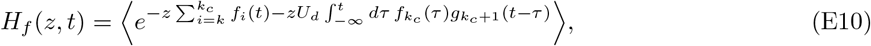

when

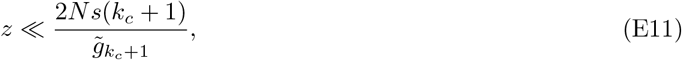

where we have used the notation from the main text, 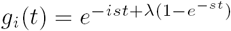.

#### The relationship between the 'feeding class' trajectory *fk*_*c*_(*t*) and the allele frequency trajectory *f*(*t*)

Equivalently, this result can be rewritten in terms of the relationship between the allele frequency trajectory *f*(*t*) and the trajectory of the portions of the alleles in classes with *k* ≤ *k*_*c*_

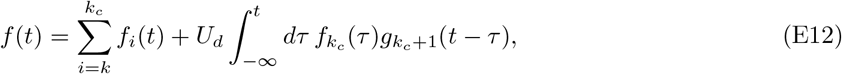

which is valid as long as 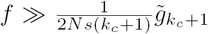. Because the expression on the right hand side of Eq. (E12) is dominated by the last term, the full allele frequency trajectory reduces to a single stochastic term *f*_*k*_*c*__. Therefore, we can calculate the distribution of *p*(*f*, *t*) near any given frequency *f* by: (1) determining the ‘feeding class’ *k*_*c*_(*f*) which corresponds to the class of lowest fitness in which genetic drift is not negligible, and (2) calculating the distribution of this time integral of the trajectory in that class, *f*_*k*_*c*__(*t*), subject to the boundary condition that 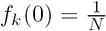.

In principle, this is still challenging if *k*_*c*_ > *k*, because the trajectory in class *k*_*c*_ still depends on the trajectories in higher-fitness classes, all of which are stochastic. In addition, calculating the distribution of the convolution of *f*_*k*_*c*__(*t*) and *gk*_*c* + 1_(*t*) is still difficult, even when *k*_*c*_ = *k*. Fortunately, a simplification arises from the highly peaked nature of *gk*_*c* + 1_(*t –* τ). Because the exponent in *gk*_*c* + 1_(*t –* τ) is peaked in time, the integral in Eq. (E12) is, up to exponentially small terms, dominated by the region in which *gk*_*c* + 1_(*t –* τ)*f*_*k*_*c*__(τ) is largest. Since the variation in the magnitude of *gk*_*c* + 1_(*t –* τ) is much larger than the variation in the magnitude of *f*_*k*_*c*__(τ), the integral will be dominated by the window during which *gk*_*c* + 1_(*t –* τ) is at its peak, as long as *f*_*k*_*c*__(τ) *≠* 0 in that window. In that case, we can make a Laplace-like approximation in Eq. (E12), in which we expand *gk*_*c* + 1_(*t –* τ) around its peak, and neglect contributions that are far away from this peak, since these are exponentially small. Near 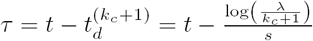,

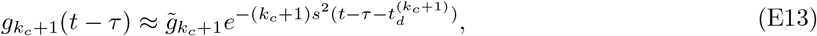

which yields

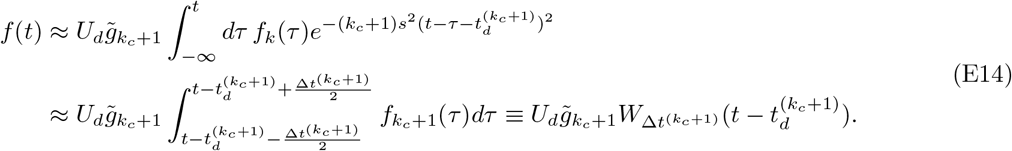

As a result of this simplification, the allele frequency does not depend on the full frequency trajectory in the feeding class *f*_*k*_*c*_+1_(*t*), but only on its time integral (‘weight’) in a window of width 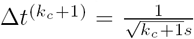 around 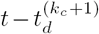, which we denote by 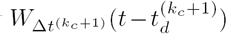. Note that Eq. (E14) implies a simple condition in terms of the allele frequency trajectory in this feeding class *k*_*c*_ that specifies when drift is negligible in downstream classes. We have shown above that as long as the total allele frequency 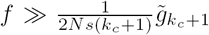, drift is negligible in classes with more than *k*_*c*_ deleterious mutations per individual. From Eq. (E14), we can see that this condition can be restated in terms of the weight in the feeding class as

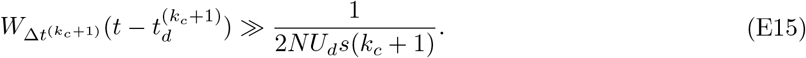

Thus, *k*_*c*_ can also be thought of as corresponding to the class of highest fitness in which the weight exceeds 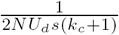.

The approximation we have used in Eq. (E14) breaks down at very early times 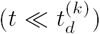, and very late times, during which *f*_*k*_*c*__(*τ*) = 0 in the relevant window. These correspond to the spreading and extinction phases of the trajectory. We show in Appendix H that the former has a negligible impact on the site frequency spectrum. The latter phase however has an important effect at very high frequencies of the mutant, i.e. when the wild type is rare and in its own extinction phase. During this extinction phase,

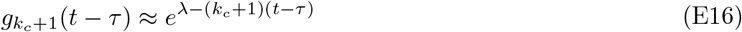

uniformly in *t* and the frequency trajectory is well approximated as

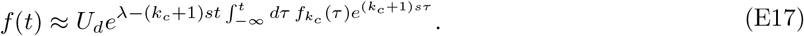

Applying the Laplace approximation once again, we conclude that the integral in Eq. (E17) is dominated by the window of width 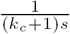 prior to extinction in the *k*_*c*_-class and therefore only weakly depends on time. Thus, during this extinction phase, the allele frequency decays exponentially at rate (*k*_*c*_ + 1)*s*, and can be written as

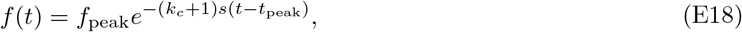

for some choice of *t*_peak_, where *f*_peak_ reflects the maximal frequency the trajectory reached before the onset of the extinction phase.

Thus, we have shown in this Appendix that the allele frequency trajectory in the peak phase of the allele only depends on the time integral of the frequency in class *k*_*c*_ over a window of specified width ∆*t*^(*k*_*c*_+1)^, and that outside this peak phase, the trajectory has an even simpler time-dependent form.

As we will see in Appendix F, the generating function for this relevant weight in class *k*_*c*_ is straightforward to calculate when *k*_*c*_ is the founding class (i.e. for *k*_*c*_ = *k*). This case is relevant for trajectories that arise in class *k* and exceed frequencies 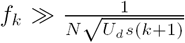, which means that the feeding class weight will exceed 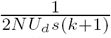 for a certain period of time.

However, not all trajectories that arise in class *k* will reach such large frequencies. We have seen in an earlier section that trajectories that do not ever exceed frequencies much larger than 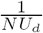 will have a trajectory that is dominated by drift throughout its lifetime. However, even those that do exceed 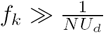, and therefore leave behind a large number of deleterious descendants will often not reach the much larger frequency 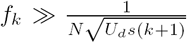. In this case, we will have to treat multiple fitness classes stochastically and the weight relevant for the peak of the trajectory will be that in class *k*_*c*_ > *k.* For *k*_*c*_ > *k*(*≥* 0), a further simplification results from the fact that the width of the window ∆*t*^(*k*_*c*_+1)^(*t*) is longer than the lifetime of the mutation in class *k*_*c*_ (see Appendix F and Appendix G for details). We use this simplification to calculate the resulting weight distribution in Appendix G. Finally, in Appendix H we use these results to obtain expressions for the average site frequency spectrum both in the case of strong and weak mutation.

### 3. Lineages on typical backgrounds 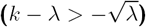

Lineages founded in classes with 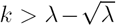 mutations will not enter the semi-deterministic regime described above. This is because selection in each individual class *i* in which they can be present prevents *f*_*i*_ from exceeding 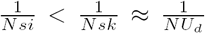, where the latter is the necessary threshold for a large enough number of deleterious descendants to be generated that their dynamics become dominated by selection in some class below the *i*-class. This threshold emerged from our analysis of the coupled branching process in the previous subsection and is further discussed in Appendix G.

In contrast to lineages arising far above the mean of the fitness distribution, the frequency trajectories of lineages that arise near the mean of the fitness distribution are dominated by drift, and eventually capped by negative selection at large enough frequencies. Selection becomes an important force about 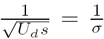 generations after the lineage was founded. At this time, the accumulated deleterious load since arising becomes large enough to impact the trajectory of the mutation. This deleterious load will impact the trajectory substantially when the frequency of the lineage *f*(*t*) becomes comparable to the ‘drift barrier’ set by its current relative fitness 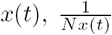. The expected fitness of a lineage founded near the mean of the distribution (with 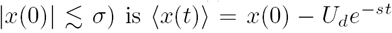. Provided that the lineage has not drifted to extinction by *t*, its expected frequency at 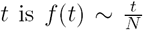. Thus, when 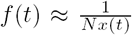, the effect of selection will dominate over drift. This occurs that 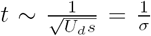. Thus, lineages that arise near the mean of the fitness distribution have a trajectory that has neutral statistics for the majority of its lifetime, but does not exceed 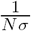. Finally, lineages arising in classes far below the mean of the fitness distribution 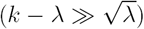, will also be dominated by drift, but limited to even lower frequencies. However, these lineages are also comparatively rare and only have a small relative impact on the lowest frequency part of the site frequency spectrum 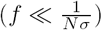.

## Appendix F: The distribution of allele frequencies and of the weight in the founding class

In this Appendix, we calculate the distribution of frequencies *f*_*k*_(*t*) and weights 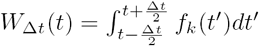 for the stochastic process defined by

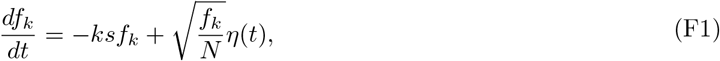

with 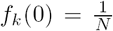 and *f*_*k*_(*t*) = 0 for *t* < 0. This process describes the trajectory of the component of the lineage that remains in the founding class (the ‘founding genotype’). To calculate these distributions, we begin by defining the joint generating function for the frequency *f*_*k*_(*t*) and the total time-integrated weight up to time *t*,

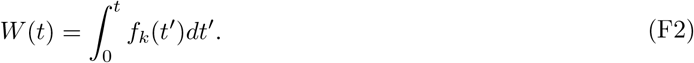

The joint generating function for these two quantities is defined as

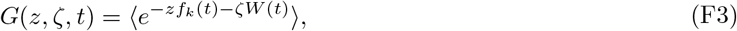

and satisfies the PDE

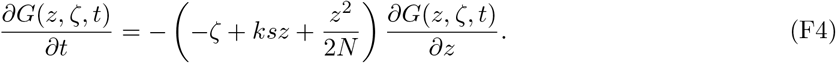

Once again, we solve this PDE using the method of characteristics. The characteristics *z*(*t−t′*) are defined by

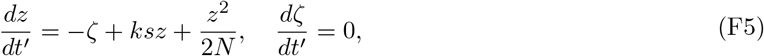

and are subject to the boundary condition *z*(*t*) = *z*, *ζ*(*t*) = *ζ.* The generating function is constant along the characteristics 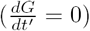, and therefore satisfies

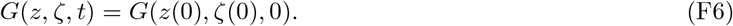

After integrating the ODEs in Eq. (F5), we find that the characteristics follow

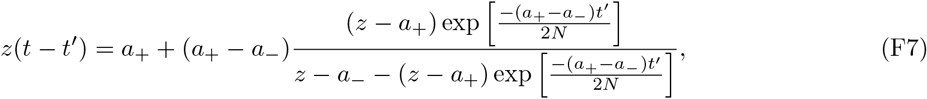

with 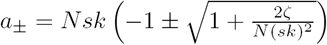.

We can verify that the correct marginal generating function for the frequency of the lineage emerges from this result by setting ζ = 0 and imposing the boundary condition 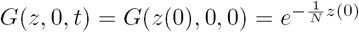, which corresponds to the initial frequency at *t* = 0 being 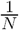.

To obtain the marginal generating function for the weight in the window between 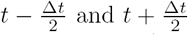, we set *z* = 0, *t* = Δt, and choose a boundary condition that reflects the distribution of frequencies 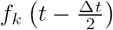 generations after the lineage was founded (see Eq. (D2)),

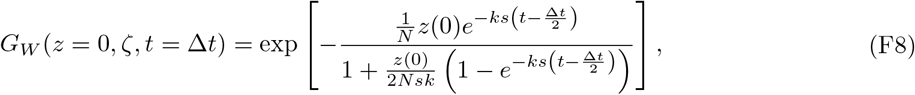

where

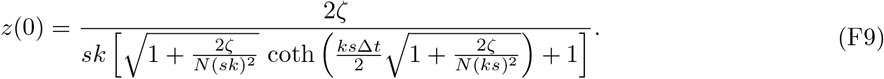

The generating function in Eq. (F8) captures the full time-dependent behavior of the weight in the founding class in a window of width Δ*t*, and can be inverted by standard methods. However, it is in practice unnecessary to invert Eq. (F8) to calculate the site frequency spectrum. For our purposes here, we will be mostly concerned with two special cases: the total weight in the founding class from founding to extinction, 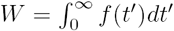, and the time-integral of the distributions of frequencies *p*(*f*(*t*)) and weights *p*(*W*_Δ*t*_(*t*)) in a window of specified width Δ*t*. The former case has been calculated previously by Weissman et al. (2009). We quote and discuss this result for completeness in the section below. We then analyze the latter case in the following section.

### 1. The distribution of the total lifetime weight in the founding class, 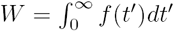

The first special case that will be relevant to our analysis of trajectories and allele frequency spectra is the total integrated weight in the founding class from founding (*t* = 0) to extinction. By setting 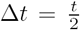 in Eq. (F8) and Eq. (F9), we find that the generating function for the total weight from founding to some later time *t* is

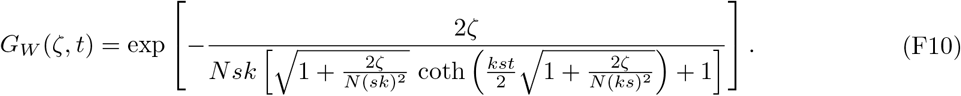

Note that Eq. (F10) becomes independent of time when 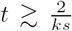 (uniformly in *ζ*), which agrees with our heuristic intuition that the lifetime of a mutation in class *k* is not longer than ~ 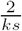 generations. Since we have shown in Appendix E that the allele frequency trajectory *f*(*t*) depends on the weight in a window of width 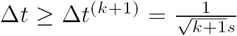 (where the ≥ sign follows because *k_c_ ≥ k*) that is longer than 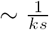 for *k* > 1 (with *k* = 1 being the marginal case), the distribution of *W*_Δ*t*(*k+*1)_ (*t*) will be either equal to the total lifetime weight of the allele (for 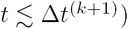) or negligible for *t* ≳ Δ*t*^(*k*+1)^.

By taking the limit *t* → ∞ in Eq. (F10), we obtain that the distribution of the lifetime weight in the founding class is

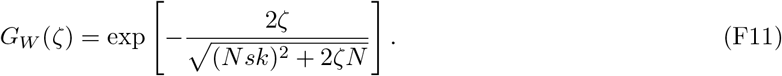

The inverse Laplace transform of Eq. (F11) can be evaluated by standard methods, which yields the distribution of the lifetime weight in the founding class

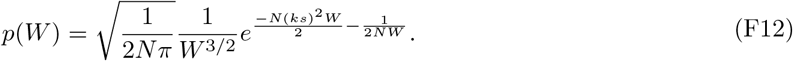

### 2. The time integrals of *p*(*f*, *t*) and *p*(*W*_Δ*t*_,*t*)

To calculate the average site frequency spectrum, we need to calculate the time-integral of the distributions of frequencies and weights over time. In principle, this can be done by inverting Eq. (F8) and then integrating the distribution of *W*_Δ*t*_(*t*) over time. However, since this is a somewhat laborious calculation, we will use a convenient mathematical shortcut in which we first solve for the distribution of weights in a different stochastic process, and then relate this back to the original process in Eq. (F1).

Specifically, we consider the stationary limit of the stochastic process defined by the Langevin equation

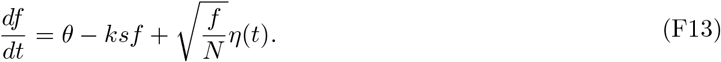

This describes the time-evolution of the frequency of a lineage with fitness –*ks* in which individuals are continuously generated by mutation at some rate *Nθ* (and have frequency 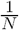 at the time when they are generated). This process is relevant because the distribution of frequencies and weights in the stationary process are related to the time-integrals of the distributions of *f*(*t*) and *W*_Δ*t*_(*t*). More precisely, in the limit that θ → 0 (keeping *N* constant), the distributions of *f* (and its time integrals) in the stationary process are the same as the time-integrated distributions of the non-stationary process, provided that we also divide by the total rate at which new individuals are generated, *Nθ*, to ensure proper normalization. That is,

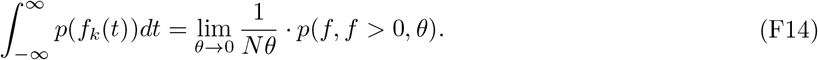

We denote the joint generating function for the frequency, *f*, and weight in this process, *W*(*t*, *θ*) = 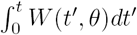, by

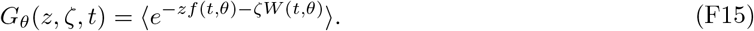

*G*_*θ*_(*z*, *ζ*, *t*) satisfies the PDE

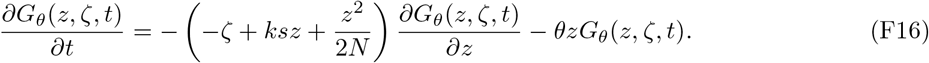

Note that the generating functions for the two processes are related, and that by setting *θ* = 0 in Eq. (F16), we obtain the generating function for the non-stationary process (see Eq. (F4)). In particular, the characteristics for Eq. (F16) are the same as the characteristics for Eq. (F4), and they follow the form we calculated previously and quoted in Eq. (F7). Along these characteristics, the generating function satisfies

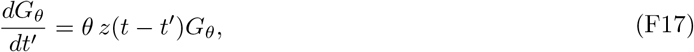

or equivalently, after integrating,

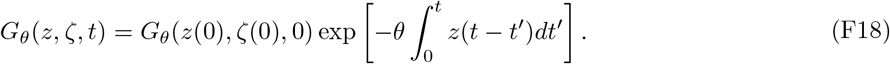

However, the boundary conditions for the two processes are different. The non-stationary process is subject to the boundary condition that there is a single individual present in the lineage at *t* = 0, *G*(*z*(0), ζ(0), 0) = 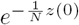, whereas the stationary process is subject to the boundary condition that the process is stationary at the initial time point, *t′* = *t.* The stationary property of the frequency distribution is guaranteed by the boundary condition 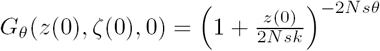. This can be obtained either by inspection, or by substituting an arbitrary boundary condition and finding the limiting form for the generating function for the frequency as *t →* ∞, and noting that *z*(0) becomes independent of *z* as *t →* ∞, so the initial condition has no impact on the frequency distribution.

Plugging in the expression for *z*(*t* – *t′*) from Eq. (F7) into Eq. (F18), and performing the integral over *t*′, we arrive at the solution to the joint generating function for *f*(*θ*) and *W*(*θ*),

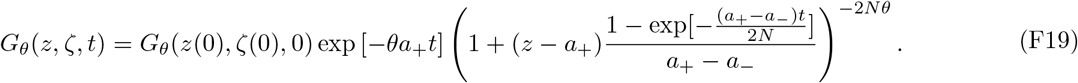

To obtain the marginal generating function for *f*(*θ*), we set ζ = 0, giving *a*_+_ = 0, a_−_ = –2*Nsk*, and

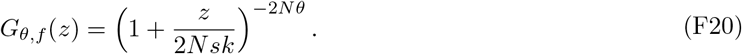

Conversely, to get the generating function for *W*(*t*, *θ*) we set *z* = 0, which after some rearranging yields

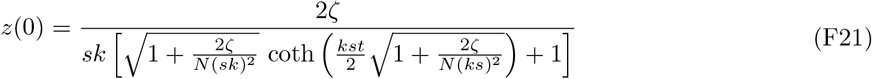

and

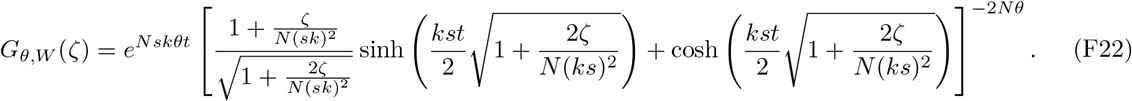

We invert Eq. (F20) and Eq. (F22) below.

#### Inversion of the generating functions in Eq. (F20) and Eq. (F22)

Since only the non-extinct portion of the process contributes to the site-frequency spectrum, when inverting the generating functions for the weight and frequency, we will use the following relationship between the probability distribution *p*(*g*) and the moment generating function *G*_*g*_(*z*) of a random variable *g:*

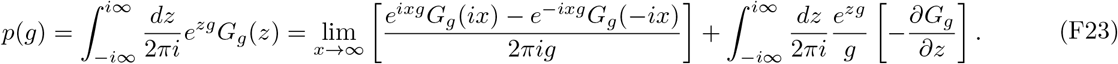

From the definition of the moment generating function and the sine limit definition of the Dirac *δ* function 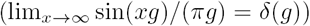, it follows that the boundary terms amount to the probability mass at *g* = 0 and that the distribution of the nonzero portion of the process is

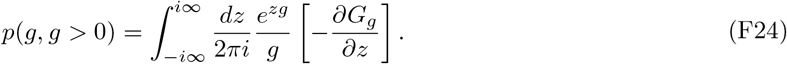

After plugging this expression and the generating function for the frequency Eq. (F20) into Eq. (F14) and taking the *θ* → 0 limit, we find that the time-integrated distribution of frequencies in the founding class is

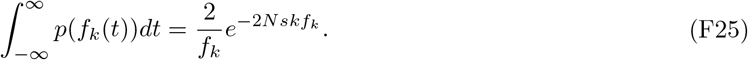

The time-integrated distribution of weights in the feeding class can be obtained in an entirely analogous fashion. In this case, it will be convenient to treat the cases *k* = 0 and *k* > 0 separately. When *k* = 0, a lengthy but straightforward substitution of Eq. (F22) into Eq. (F24) gives

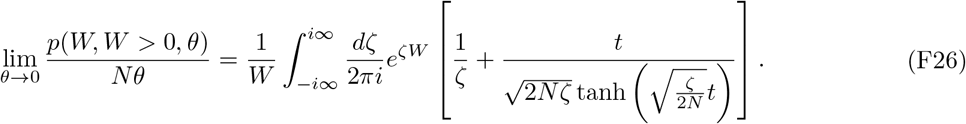

The simplest way to carry out this integral is by contour integration. To do this, we close the contour using a large semi-circle in the left half-plane. The contribution from this circle vanishes as the radius of the semicircle approaches infinity, and so the integral considered above is equal to the sum of the residues within the left half-plane. The integrand has simple poles at 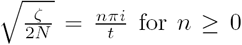 with residues 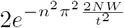, which yields

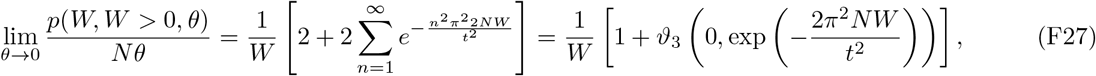

where *ϑ*_3_ is the elliptic theta function. Asymptotic expansions for small and large arguments give

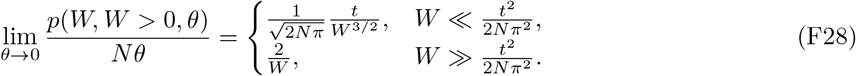

The case *k* > 0 is slightly more straightforward to evaluate, since the length of the intervals we are interested in is longer than the typical timescale of selection 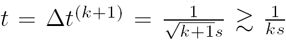. As a result, the arguments in the hyperbolic functions in Eq. (F22) satisfy 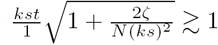 (for *k* > 1; with *k* = 1 being the marginal case), which yields a simple form for the distribution of nonzero weights

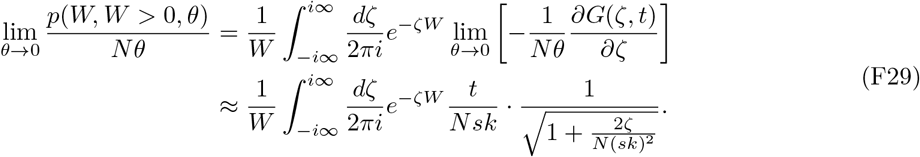

Note that the expression in Eq. (F29) reduces to a standard Gaussian integral. By carrying out this integral, we obtain for the time integral of the distribution of weights in the founding class

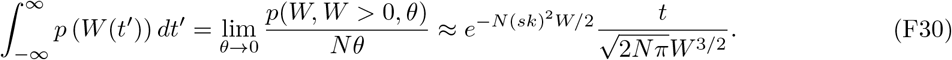

## Appendix G: The distribution of weights in classes below the founding class

We have seen in Appendix E that when the allele frequency trajectory in the founding class *f*_*k*_(*t*) is small enough, the effects of genetic drift cannot be ignored in multiple fitness classes. In this section, we consider how the trajectories (and their weights) in these stochastic classes are coupled, and derive the distribution of lifetime weights in class *k +* Δ, in which individuals carry Δ more mutations compared to individuals in the founding class.

### 1. The relationship between the trajectory in the founding class *k*, and the weight in class *k +* 1

We begin by considering the total lifetime weight in the class right below the founding class (*i* = *k +* 1), which we will denote *W*_*k*+1_. *W*_*k*+1_ clearly depends on the weight in the founding class, *W*_*k*_, since the total number of mutational events from the *k*-class into the *k+*1-class is equal to *NU*_*d*_*W*_*k*_. As we describe in Appendix B, each one of these mutational events founds a ‘sub-lineage’ and the stochastic trajectory of each sub-lineage is described by Eq. (F1). The total weight of the lineage in class *k+*1 is simply the sum of the weights of each of these sub-lineages. The generating function of the lifetime weight in the *k*+1 class, *W*_*k*+1_, is related to the lifetime weight in the founding class *W*_*k*_ according to

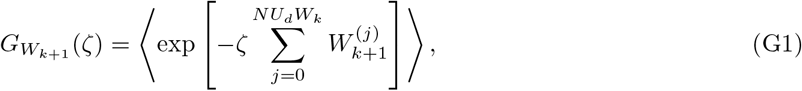

where 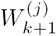 denotes the weight of the sub-lineage founded by the *j*^th^ mutational event. Since the 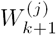 are independent and identically distributed, the generating function of their sum is equal to the product of their generating functions and

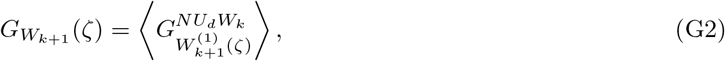

where the final average is taken over the distribution of the weight, *W*_*k*_, in the *k*-class. The generating functions of *W*_*k*_ and 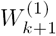 are both given by Eq. (F11).

Using the same methods that we used to invert Eq. (F11), we obtain that the distribution of the total weights in class *k* + 1, conditioned on the weight in class *k* being equal to *W*_*k*_, is

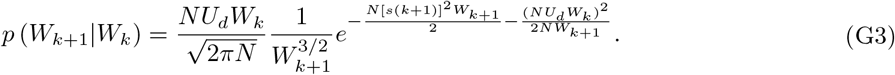

We can see from this equation that the neutral decay of the distribution of weights in class *k* + 1, which results from drift and is proportional to 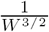, is exponentially cut off for 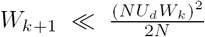 and for 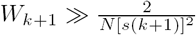. The latter, high-weight cutoff is familiar from before, and results from selection within the *k* + 1 class. The low-weight cutoff results from the pressure of incoming mutational events.

A simple heuristic can explain the dependence of the low-weight cutoff on the weight in the founding class, *W*_*k*_. The weight *W*_*k*+1_ is at least as large as the weight of the largest sub-lineage. Because each of the *NU*_*d*_ *W*_*k*_ mutational events generates a sub-lineage that survives for *T* generations with probability 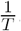, and leaves a weight of order 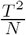, at least one of these sub-lineages will survive for *T* generations with probability equal to 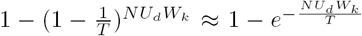. This probability is of order 1 for *T* ~ *NU*_*d*_*W*_*k*_, which means that with probability order 1 at least one of the sub-lineages will have weig 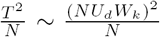. Note that this also means that when 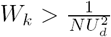 (consistent with the lineage exceeding frequency 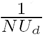 in the founding class), the weight in the next class is guaranteed to be larger than the weight in the founding class. This means that lineages that exceed the frequency 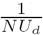 in the founding class are almost guaranteed to generate an even larger number of individuals in the next class, which generates an even larger number of individuals in the following class, and so on.

We have implicitly assumed that the trajectory of each of the sub-lineages is dominated by drift. This will be true as long as 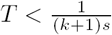 (i.e. as long as 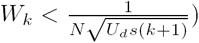. In contrast, when 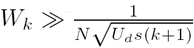, a large number the lineages will exceed the frequency 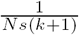 in the next class, and the trajectory in that class will become dominated by selection. We have shown in Appendix E that once this happens, drift in class *k +* 1 and all classes below it will become negligible. Note that this heuristic argument also explains the self-consistency condition that emerged in Appendix E (see Eq. (E15)), and explains why genetic drift becomes neglibile in the *k*_*c*_ + 1 class whenever the weight in the *k*_*c*_-class is larger than 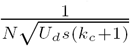.

In the section below, we will use the insights above to evaluate the weight distribution in class *k +* Δ, conditioned on the lineage arising in class *k* and selection being negligible in all classes beneath it, *W*_*i*_ ≪ 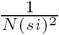 for *i* ≤ *k* + Δ. Because the lifetime of the longest-lived sub-lineage in each of these classes is at most 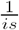 in this limit, and because the sub-lineages are seeded into the *i*-class over a time that is, by assumption, shorter than 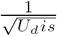, the total lifetime of the lineage in all of these classes is strictly shorter than Δ*t*^(*i*)^, which is why we do not need to be concerned with the full, time-dependent properties of the distribution of weights in this class. Instead, the calculation of the distribution of lifetime weights will suffice for calculating the site frequency spectrum.

### 2. The distribution of the weight in class *k* + Δ

Having obtained the distribution of the weight in class *k +* 1, conditioned on the weight in class *k* being equal to *W*_*k*_ (see Eq. (G3)), we can calculate the marginal distribution of weights *W*_*k*+1_ by averaging over *W*_*k*_. In the limit that 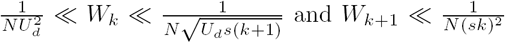 that we are interested in here, this distribution is

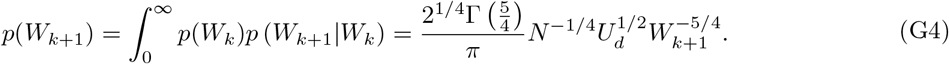

Note that the distribution in the (*k +* 1)-class decays less rapidly than in the *k*-class. In particular, the probability that the weight in the (*k* + 1)-class exceeds 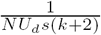 (and leads to the deterministic propagation of individuals in classes with *k +* 2 or more deleterious mutations) is

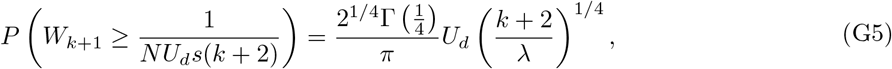

which is larger than the probability that the weight in the *k*-class exceeds the corresponding value by a large factor ~ *λ*^1/4^, consistent with our intuition that the weight in the class below the founding class is guaranteed to exceed the weight in the founding class if 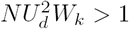.

In general, we can calculate the distribution of the weight in class *k* + Δ by iterating this procedure. Specifically, the distribution of the weight in class *W*_*k*+2_ conditioned on the weight in class *k*+1 being equal to *W*_*k*+1_ also follows Eq. (G3) (but with *k* changed to *k +* 1). By repeating the above procedure Δ times, we find that the distribution of lifetime weights in class *k +* Δ is

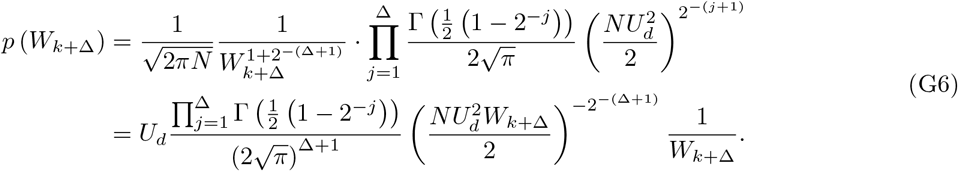

## Appendix H: The site frequency spectrum

In this Appendix, we use the results obtained in previous sections to calculate the site frequency spectrum of the labelled lineage in the limits that *f* ≪ 1 and 1 – *f* ≪ 1, by evaluating and inverting the generating function *H*_*f*_(*z*, *t*) for the total frequency of the labelled lineage.

### 1. Weak mutation (*U*_*d*_ ≪ *s*)

We have seen in Appendix D that trajectories of mutations in the presence of weak background selection (*U*_*d*_ ≪ *s*) are to leading order in the small parameter *λ* same as those of unlinked loci with fitness −*ks.* In Appendix F we have shown that the time-integrated distribution of allele frequencies of a single unlinked locus of fitness –*ks* is

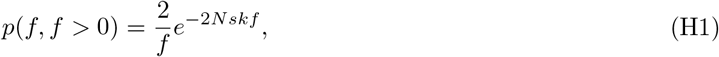

which agrees with classical results by Ewens (1963) and Sawyer and Hartl (1992). Thus, the contribution to the site frequency spectrum of neutral mutations arising in class *k* is

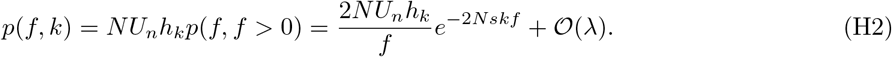

Summing the contributions of all the classes, we find that the full neutral site frequency spectrum is

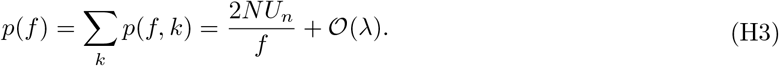

The site-frequency spectrum of deleterious mutations follows from the same argument, since the trajectory of a deleterious mutation arising on the background of an individual with *k* deleterious mutations is the same as the frequency trajectory of a neutral mutation arising in an individual with *k + 1* deleterious mutations. Thus, the site frequency spectrum of deleterious mutations is

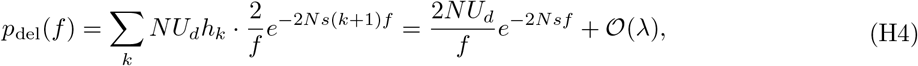

which once again agrees to leading order with the site frequency spectrum that we would have obtained assuming that all selected sites at the locus were unlinked.

### 2. Strong mutation (*U_d_* ≫ *s*)

In the presence of strong mutation, we have seen that trajectories of mutations are dominated by drift at the lowest frequencies, where the generating function reduces to the generating function of a neutral mutation, and is simply equal to the *k* = 0 limit of the single locus generating function in Eq. (D2). We have already calculated the site frequency spectrum that results from these trajectories in the previous section. Plugging in these results, we find that

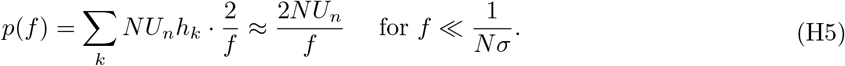

The site frequency spectrum at these frequencies is dominated by the contributions of lineages arising in average backgrounds, with 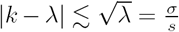. By the same argument, the frequency spectrum of deleterious mutations at the same frequencies is also

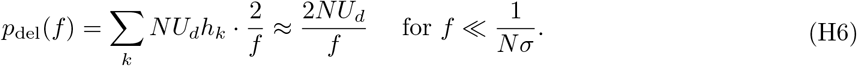

At larger frequencies, the site frequency spectrum becomes dominated by lineages arising in unusually fit backgrounds, with 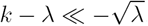. Their trajectories are instead described by Eq. (E10). We have seen that the integral in the exponent of Eq. (E10) has a different dependence on 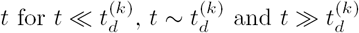, which we have labelled the 'spreading', 'peak' and 'extinction' phases of the trajectory. In evaluating the site frequency spectrum *p*(*f*), it will be convenient to calculate the contributions from each of these phases separately. We denote these contributions as *p*_spread_ (*f*), *P*_peak_(*f*) and *p*_ext_(*f*), and the full site frequency spectrum is obtained by summing,

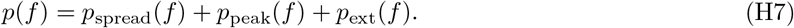

We evaluate *p*_peak_(*f*) and *p*_ext_(*f*) in the next two subsections of this Appendix. Then we show in the last subsection of this Appendix that the contribution from *p*_spread_(*f*) is sub-dominant to that of *p*_ext_(*f*).

#### a. Contribution from the peaks of trajectories

In Appendix E, we have shown that in the peak phase of the trajectory, the total allele frequency is

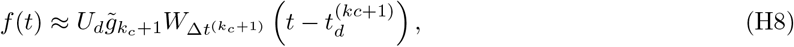

where *k*_*c*_ is the class with the largest number of mutations for which 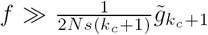, or equivalently, the class with the smallest number of mutations in which the weight exceeds 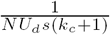.

We have seen above in Appendix G that to achieve such a large weight in class *k*_*c*_, a mutation could have arisen in class *k* = *k*_*c*_ and traced an unusually large trajectory, or arisen in class *k_c_ −* 1, and traced a smaller trajectory in that class, which led to the creation of a large number of deleterious descendants in class *k*_*c*_, at least one of which had weight exceeding 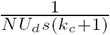. Alternatively, it could have also arisen in class *k_c_ −* 2 and traced an even smaller trajectory in that class, that led to a larger weight in class *k_c_ −* 1, and a sufficiently large weight in class *k*_*c*_ for genetic drift to be negligible in classes *i* > *k*_*c*_ + 1. In other words, in the range of frequencies

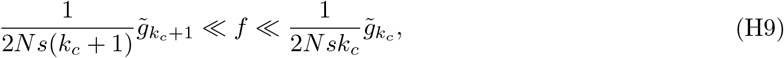

we see the peaks of trajectories originating in classes *k* < *k*_*c*_, as long their weight in class *k*_*c*_ is large enough that genetic drift in classes of lower fitness can be ignored. All of these peaks contribute to the site frequency spectrum and by integrating Eq. (H8) in time, we find that

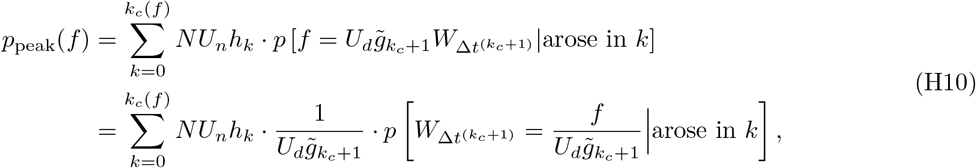

where the last term represents the time-integrated distribution of weights in a window of width 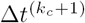 in class *k*_*c*_ of a lineage that arose in class *k.* This distribution is given by Eq. (F28) for *k*_*c*_ = 0. Otherwise, when *k*_*c*_ > 1, the time-integrated distribution in Eq. (H10) is equal to the product of the window width, Δ*t*^(*k_c_*+1)^, and the distribution of lifetime weights in the founding class, given in Eq. (G6).

Since we have previously calculated all of these quantities, we can now turn to evaluating the sum in Eq. (H10). When 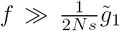, then *k*_*c*_ = 0, and the sum in Eq. (H10) has only one term (*k* = 0). By substituting in the expression for the time-integrated distribution of weights in Eq. (F28), we find that

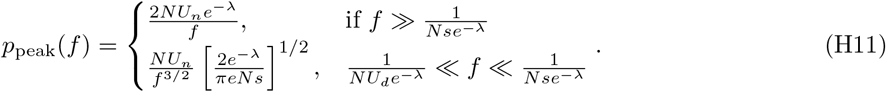

At lower frequencies, 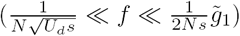, lineages originating in multiple different fitness classes will be able to contribute to the site frequency spectrum. At these frequencies,

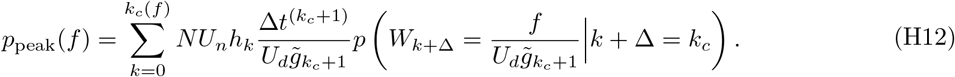

Plugging in the expression for *p*(*W*_*k* + Δ_) from Eq. (G6), we find

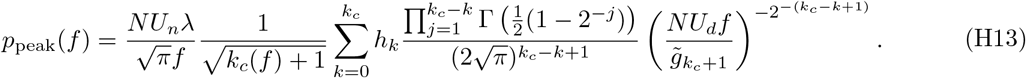

Because *λ* ≫ 1, this sum is dominated by the *k* = *k*_*c*_ term, as *h*_*k*_ decays much more rapidly with decreasing *k* than any of the other terms increase. To evaluate the *f*-dependence of this term for 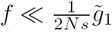 and *k*_*c*_(*f*) ≫ 1, we can solve the self-consistency condition for 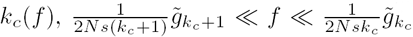, by setting 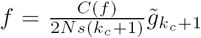 for some constant 1 ≪ *C*(*f*) ≪ *λ*. By solving for *k*_*c*_(*f*), we find that to leading order the relevant fitness class is

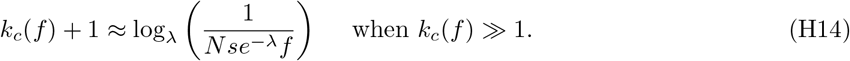

Plugging in, we obtain that the leading order term in the distribution of peak sizes is

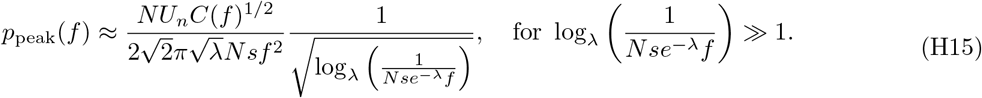

The term *C*(*f*) depends on *f* weaker than logarithmically, and on frequency scales on which *p*_peak_(*f*) changes substantially it will be approximately constant, *C*(*f*) ≈ *C.*

Because the crossover between the *f*^− 3/2^ scaling of *p*_peak_(*f*), which occurs at high frequencies 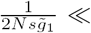 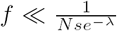 (where *k*_*c*_(*f*) + 1 = 1), and the 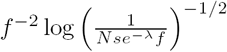 behavior, which is valid at substantially lower frequencies (where *k*_*c*_(*f*) ≫ 1), is in principle broad, this constant factor *C* is difficult to determine: asymptotic matching does not typically work well in the presence of such broad transitions, and crude ‘patching’ methods do not, in general, offer satisfactory results (Hinch, 1991). Thus, Eq. (H15) is undetermined up to the constant factor *C*^1/2^, which is between 1 and 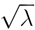. For our purposes here, this level of precision is sufficient — 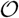(1) precision in the form of the spectrum was, after all, expected in the Laplace-like approximation that we used in Appendix E to calculate the stochastic integral over the trajectory of the feeding class. Thus, by absorbing 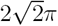 term into this constant factor, and relabeling *C*^1/2^ as *C*, we find that the peak contribution to the site frequency spectrum is

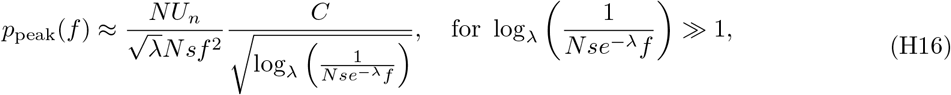

with *C* in the range 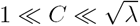.

#### b. Contribution from the extinction stage of trajectories

Once the trajectory is beyond its peak, the total allele frequency decays as

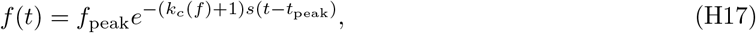

where *f*_peak_ denotes the maximal frequency that the trajectory reaches and Eq. H17 is valid for 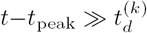. Note that this stage only exists for frequencies 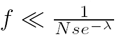. At higher frequencies, 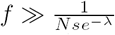, the total allele frequency simply mirrors smoothed fluctuations in the founding class. Eq. (H17) can be straightforwardly integrated in time to obtain the contribution of this trajectory to the site frequency spectrum

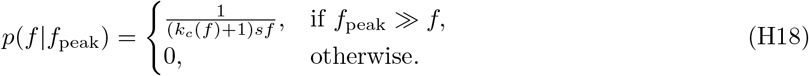

Averaging Eq. (H18) over all possible trajectories, we find that

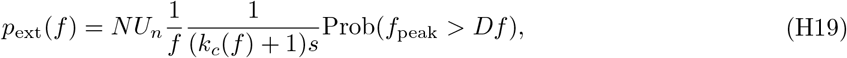

where *D* > 1 is a constant that we have introduced to correctly account for the fact that the peak phase occurs at frequencies that are at least 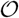(1) higher than the frequencies in the extinction stage.

For peak frequencies 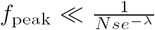, we have already calculated the overall time-integrated distribution of peak sizes of lineages arising in classes of all fitness, and we can use this result to calculate the total probability that a trajectory passes through *f* in its extinction stage,

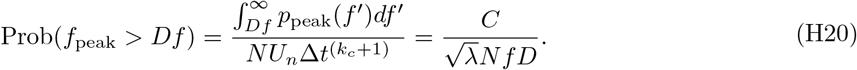

This means that the contribution to the site frequency spectrum from the extinction phase of trajectories is equal to

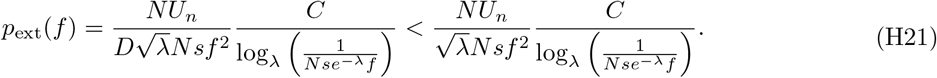

Therefore, *p*_ext_(*f*) is strictly smaller than *p*_peak_(*f*) by a factor 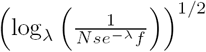, which is large when 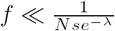. Thus, this phase of the trajectory has a small effect on the low-frequency end of the spectrum. However, in the high frequency end of the spectrum, when 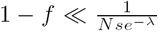, the only contribution comes from this ‘extinction’ phase of the wild-type, which starts once the mutant approaches the frequency *f* ≫ 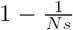 in the 0-class. These events happen at rate equal to 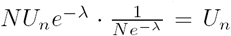, and each contributes 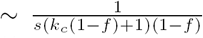 to the site frequency spectrum. Multiplying these two terms, we find that the site frequency spectrum is proportional to

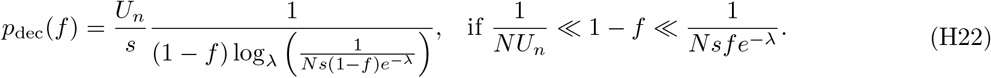

#### c. Contribution from the spreading stage of trajectories

At frequencies 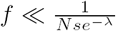, the site frequency also receives contribution from the spreading stage of trajectories, in which the allele frequency rapidly increases as the allele spreads through the fitness distribution. In this stage, the rate at which the frequency increases is strictly larger than what it would be if we ignored any contributions from the founding class after the mutation exceeds frequency 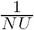, (i.e. assuming 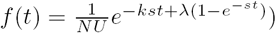,

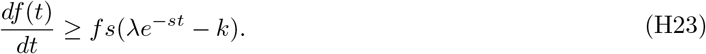

Far below the peak of the trajectory, where *λ*e^*−st*^ ≫ *k*, the contribution from this stage of a single trajectory to the frequency spectrum that passes through *f* is thus simply bounded by

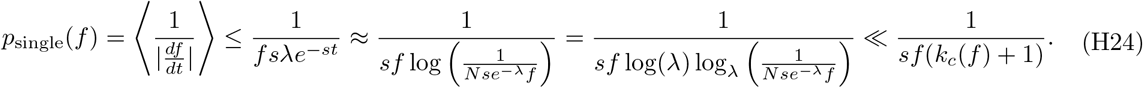

Since the number of trajectories that pass through frequency *f* in the spreading phase is the same number that pass through *f* in the extinction phase, the contribution from the spreading phase to the site frequency is strictly smaller than that of the extinction phase throughout the region where both contributions exist, 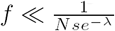.

## Appendix I: Distributions of effect sizes

When the effects of deleterious mutations are not all identical, but instead have a distribution with finite width, *ρ*(*s*), the deterministic dynamics that arise through the combined action of mutation and selection will be modified. In this Appendix, we consider these deterministic dynamics. For concreteness, we assume that the fitness effects of new mutations come from a gamma distribution with mean *s* and shape parameter *α*,

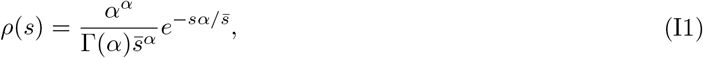

and that these deleterious mutations occur at an overall rate *U*_*d*_.

Under the assumption that all mutations have strong enough effects on fitness that the fitness of the population at the locus does not experience Muller’s ratchet on timescales of coalescence, the mean fitness of an allele at the locus will be equal to –*U*_*d*_, with the most-fit individuals being those with no deleterious mutations and an absolute fitness equal to 0. Consider now the deterministic dynamics of a lineage founded in an individual at absolute fitness –*x.* The fitness of the lineage founded by this lineage will change as it accumulates new deleterious mutations according to

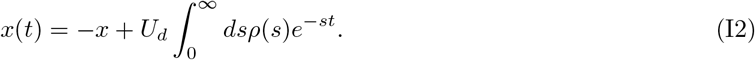

Evaluating this integral, we find

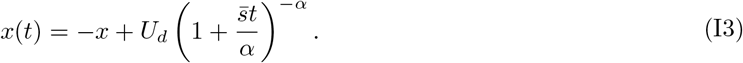

When *α* is sufficiently large, corresponding to a sufficiently narrow fitness distribution, the resulting trajectory is well approximated by assuming that all fitness effects are the same and equal to the average fitness 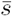 (or, more precisely, the harmonic mean of 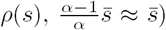. To calculate how large *α* needs to be for this approximation to be valid, we can calculate the determinstic expectation for the average number of individuals in the lineage at time *t* after founding. This quantity is equal to

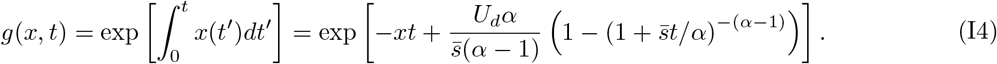

We see that this differs from the single-s expression only in the last term, proportional to 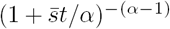. At sufficiently short times, 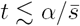, this is well-approximated by 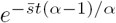. On sufficiently long timescales, this will not be the case. However, because the overall magnitude of this term becomes negligible at times long after the peak of *g*(*x*, *t*), *t* ≫ *t*_*d*_, we only need it to remain well-approximated by an exponential on timescales *t* ≲ *t*_*d*_, which requires that 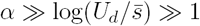. When this is the case, *g*(*x*, *t*) is, up to perturbative corrections, given by

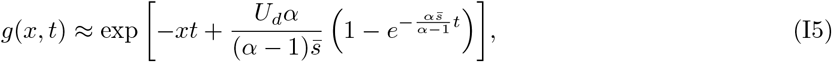

and the effects of selection are well-described by a single-s model on all timescales.

